# Generation mechanisms and species-specific properties of parkinsonian beta oscillations in the basal ganglia

**DOI:** 10.64898/2026.09.24.753820

**Authors:** Elena Nicollin, Marc Deffains, Nicolas P. Mallet, Arthur Leblois

## Abstract

The basal ganglia (BG), thalamus and cortex form the BG–thalamo–cortical (BGTC) network, which is essential for voluntary movement and centrally involved in Parkinson’s disease (PD). In both patients and animal models, BG neuronal activity exhibits exaggerated oscillatory synchronization in the beta frequencies (13–30 Hz). Theoretical studies have proposed multiple network mechanisms for the generation of these abnormal beta oscillations. However, key properties, such as frequency and power distribution across BG nuclei, vary substantially among patients and between animal models (rodents vs. non-human primates). This variability complicates direct comparisons between theoretical models and experimental data, and questions whether distinct neuronal mechanisms may underlie beta oscillations across species.

In an experimentally constrained BGTC network model with species-specific parameters (synaptic, neuronal, and network properties), we evaluate the features of abnormal beta oscillatory activity generated by different mechanisms. The network’s negative feedback loops serve as potential sources of spontaneous oscillations. Using rodent- or primate-constrained models, we compare the spectral properties of oscillatory activity across network populations for each loop and derive the expected phase relationships between populations to align predictions with existing rodent data and propose testable hypotheses for primates. We also demonstrate how oscillation frequency can be modulated when multiple generation mechanisms interact as coupled oscillators in the full network. Our results, combined with observed cross-species beta oscillations characteristics, suggest that abnormal beta oscillations likely arise from distinct mechanisms in rodents and primates.

**Significance statement:** A hallmark of Parkinson’s disease is exaggerated beta-frequency (13–30 Hz) oscillatory synchronization in the basal ganglia–thalamo–cortical network. Yet beta oscillation properties vary greatly between patients and animal models, and whether they arise from a common mechanism across species remains unknown. Using an experimentally constrained computational model with species-specific parameters, we identify which circuit may generate beta oscillations, predicting their frequency, power distribution, and phase signatures. Our results suggest that abnormal beta oscillations likely arise from distinct mechanisms in rodents and primates, and yield testable hypotheses for primates. Moreover, we show that interacting mechanisms can jointly modulate oscillation frequency. These findings offer experimentally verifiable predictions to guide future research and refine how preclinical findings translate to patients.

## Introduction

The basal ganglia (BG) are a set of subcortical nuclei interacting with the thalamus and cortex, involved in various functions, including voluntary motor control (Alexander et al., 1986). Dysfunction of the BG-thalamo-cortical (BGTC) network underlies movement disorders such as Parkinson’s disease (PD). Oscillatory synchronization in the beta frequency range (13–30 Hz) is present across multiple nodes of the BGTC network in normal and pathological states. Under normal conditions, beta activity appears during sensorimotor and cognitive tasks as brief, transient bursts at ∼20 Hz across the BGTC network (Courtemanche et al., 2003; Leventhal et al., 2012; Khanna and Carmena, 2015). In PD, however, beta synchronization becomes excessive and persistent. In experimental PD models, abnormal beta-band synchronization emerges following dopamine depletion within and between BG nuclei, BG-recipient thalamus, and motor cortex, both in non-human primates (NHPs) (Nini et al., 1995; Raz et al., 2001; Deffains and Bergman, 2019) and rats (Magill et al., 2001; Brazhnik et al., 2016; Sharott et al., 2017). In PD patients, intraoperative recordings during deep brain stimulation (DBS) surgery reveal pronounced beta oscillations in the subthalamic nucleus (STN) (Levy et al., 2000; Amirnovin et al., 2004; Kühn et al., 2005) and the internal globus pallidus (GPi) (Levy et al., 2001; Silberstein et al., 2003). These oscillations are coherent across the STN, pallidum, and sensorimotor cortex (Brown et al., 2001; Marsden et al., 2001) and are absent or less prominent in other neurological conditions, underscoring their pathophysiological specificity (Munhoz et al., 2026). While STN beta power correlates with akinesia and rigidity (Kühn et al., 2006; Sharott et al., 2014; Neumann et al., 2016) and decreases post-treatment (Wingeier et al., 2006; Kühn et al., 2008), it is unlikely to directly cause motor symptoms, as these precede beta emergence during dopamine depletion (Leblois et al., 2007; Degos et al., 2009; Quiroga-Varela et al., 2013). Nevertheless, beta power remains a robust biomarker for DBS electrode implantation and adaptative strategies (Little and Brown, 2020; Tinkhauser et al., 2020). The neural mechanisms underlying pathological beta oscillations remain largely unknown.

Absent in vitro, beta oscillations likely emerge from interactions among BGTC neuronal populations (Boraud et al., 2005). From a theoretical perspective, delayed negative feedback loops represent a minimal requirement for the spontaneous generation of oscillations in multi-population networks (Ermentrout et al., 2001). The BGTC network contains multiple such loops, with proposed candidate circuits including the subthalamo-pallidal loop, hyperdirect cortico-STN-GPi-thalamocortical pathway, striato-pallidal circuits, and intrinsic cortical/striatal generators (Pavlides et al., 2015; Rubin, 2017). Structural and physiological properties of the BGTC network shape these loops’ dynamics and constrain achievable oscillation frequencies (Zang et al., 2024). While these mechanisms have been extensively studied in isolation, how their interactions shape pathological beta remains poorly understood. Recent evidence suggests that beta oscillations in parkinsonian rats may arise from the interaction between striato-pallidal loops and pallidal recurrent inhibition (Azizpour Lindi et al., 2024). A further complication arises from species differences in beta activity. In rats, abnormal oscillations are typically 15–30 Hz (“high-beta”) (Mallet et al., 2008b; Avila et al., 2010), while in NHPs, they occur at 11–15 Hz, overlapping with alpha-band (Wichmann and DeLong, 2003; Deffains and Bergman, 2019). Furthermore, distinct circuit mechanisms appear to contribute to beta generation: while the STN is necessary for beta oscillations in parkinsonian NHPs (Tachibana et al., 2011), its inhibition does not suppress beta activity in parkinsonian rats (De la Crompe et al., 2020).

Here, we examine single candidate oscillation-generating circuits and their interactions in biologically inspired computational models of rodent and primate BGTC networks including all loops previously hypothesized to drive beta. While species-specific frequency ranges constrain plausible mechanisms for the pathological beta, we propose that, beyond frequency, phase relationships between coherently oscillating populations may be key to identifying these mechanisms.

## Materials & Methods

### Model architecture

Our model of the BGTC network includes the major populations of neurons and synaptic connections involved in negative feedback loops, resulting in a model with nine populations of neurons (Fig 1). The input structure to the BG network is represented by the primary motor cortex (Ctx). The BG itself contains the subthalamic nucleus (STN), as well as three subpopulations of the striatum (D1, D2 and FSI). The external segment of the globus pallidus (GPe) is divided into two populations to reflect its dichotomous organization into prototypical (Proto) and arkypallidal (Arky) neurons. It has been shown in rats that these two subpopulations differ functionally and anatomically (Mallet et al., 2012; Aristieta et al., 2021). The high-frequency discharge prototypic neurons promote locomotion and send projections to the striatum, the STN and the output structures of the BG. Meanwhile, the low-frequency discharge arkypallidal neurons inhibit locomotion and only innervate the striatum. The collaterals among the GPe are also asymmetrical, with prototypic cells inhibiting arkypallidal cells. While this same dichotomy remains to be established in primates, studies have shown the existence of neuronal subpopulations in the GPe (Katabi et al., 2023), which could be homologous to the Proto and Arky subpopulations. The GPi and the substantia nigra pars reticulata (SNr), despite acting on different functional systems, usually work together in parallel, receiving the same inputs from the BG and both inhibiting the thalamus (Parent and Hazrati, 1995a). We combined them into a single output population in our model, which we simply refer to as the GPi. Functional studies on NHP usually analyze GPi activity while rodent studies focus on both SNr and GPi: we therefore used GPi data whenever possible, but also resorted to using SNr data when missing parameters for GPi. Finally, the ninth population included in the model is the thalamus.

**Figure 1.**
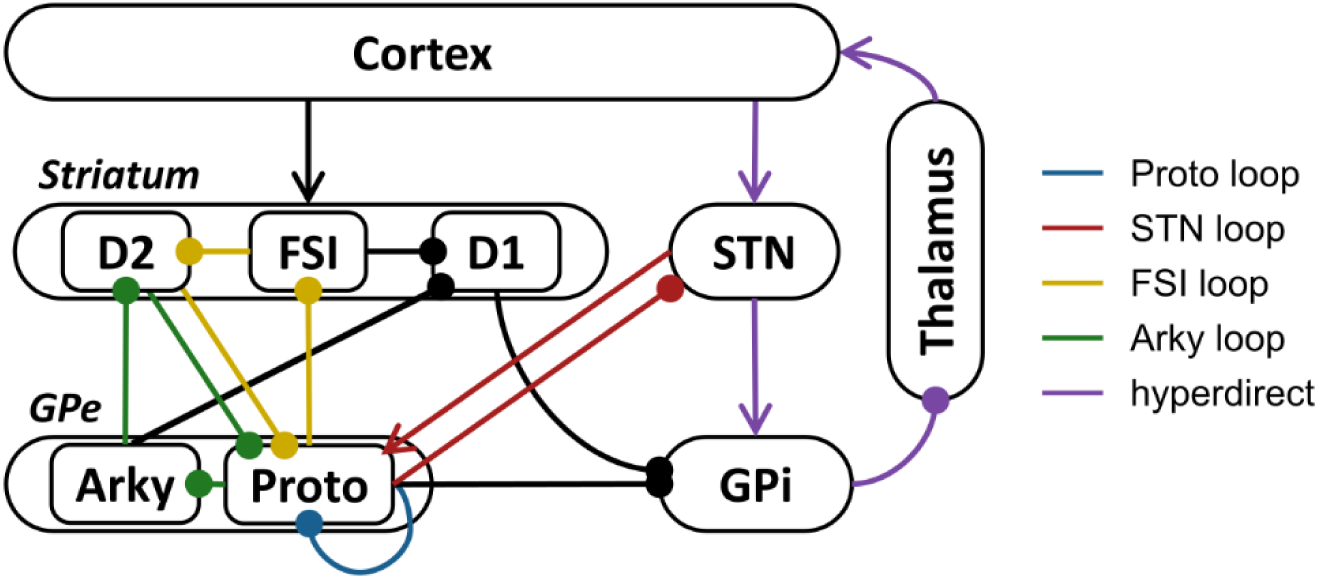
Architecture of the full BGTC network for simulation. Synaptic projections (arrows = excitatory, circles = inhibitory) are color-coded to show five negative feedback loops of interest in the generation of oscillations. Arky: arkypallidal neurons; FSI: fast-spiking interneurons; GPe: external globus pallidus; GPi: internal globus pallidus; Proto: prototypic neurons; STN: subthalamic nucleus.

Regarding the connectivity between these populations, we retained the main projections across the BGTC network, starting with all connections pertaining to known negative feedback loops. Our model contains five such loops: the shortest one corresponds to the lateral inhibition among the Proto neurons (Sims et al., 2008; Ketzef and Silberberg, 2021) and is referred to in this study as the Proto loop. The recurrent loop composed of the STN and the Proto populations, involving excitatory inputs from the first to the second (Kita and Kitai, 1991; Parent and Hazrati, 1995b) and inhibitory inputs in the opposite direction (Kita et al., 1983; Parent and Hazrati, 1995b), is referred to as the STN loop. Two loops involving the striatum and the GPe, comprised only of inhibitory projections, share as a common edge the inputs from D2 to Proto (Kita, 1994; Goldberg and Bergman, 2011). One loop then involves the upstream projection from Proto to FSI (Bevan et al., 1998; Sato et al., 2000), which then project back to D2 (Mallet et al., 2005): this loop is referred to here as the FSI loop. The second loop involves the GPe collaterals from Proto to Arky neurons (Aristieta et al., 2021; Ketzef and Silberberg, 2021), which in turn project to the D2 (Mallet et al., 2012; Fujiyama et al., 2016) and is referred to here as the Arky loop. Finally, the largest loop in the model follows the hyperdirect pathway: cortical excitatory inputs to the STN (Kitai and Deniau, 1981; Parent and Hazrati, 1995b) are relayed to the GPi (Van Der Kooy and Hattori, 1980; Parent and Hazrati, 1995b), which sends inhibitory connections to the thalamus (Clavier et al., 1976; Ilinsky et al., 1985), which in turn projects back to the Ctx (Ilinsky et al., 1985): we refer to this as the hyperdirect loop.

The proper modeling of the negative feedback loops made the implementation of these synaptic connections crucial; however, we also selected a few other important projections to include in our model. We decided to conserve the key components of the classical direct and indirect pathways: as such, we ensured cortical inputs, aside from targeting the STN, also targeted all three striatal populations (Parent and Hazrati, 1995a; Kincaid et al., 1998): besides reproducing motor pathways, these inputs also allow us to simulate the effects of cortical electrical stimulation in the downstream nuclei of the network. Secondly, the FSI lateral inhibition of D2 neurons was extended to D1 neurons (Mallet et al., 2005), and the Arky inhibition onto D2 was also generalized to D1 neurons (Sato et al., 2000; Mallet et al., 2012). Lastly, we included the inhibition of the GPi by the Proto neurons (Hazrati et al., 1990; Kincaid et al., 1991), corresponding to the output signal of the indirect pathway.

We refrained from including more connections for three main reasons. The first reason is that the dense interconnectedness of the BGTC network makes it particularly tricky to draw a clear line between relevant and irrelevant synaptic connections: we therefore settled for the minimal network projections necessary for the two objectives of our model, which are the generation of beta oscillations and the robust transfer of cortical information to the output nuclei. The second reason is that we establish the parameters of the model based on the electrophysiological and anatomical data in the existing literature, which is already unavailable for some of the connections in our model, and is likely even less so for newer, less characterized projections. The last reason is that we use the same architecture to model the BGTC network of two different species, and we cannot reliably assume that our model would remain robust for both species if it were to include a greater number of connections.

The network size was scaled down to reduce computational costs, using a fixed number of neurons in each of the simulated populations (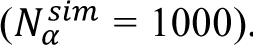). While neuron number is widely different between the various populations of the network in all vertebrates (with lower numbers of neurons in BG output populations), we deliberately choose the same number of neurons in each populations as reproducing accurately the neuronal dynamics of a large neuronal network requires minimal neuron numbers in single populations (>100 for rate model, >1000 for spiking networks, see Brunel (2000)), and increasing the population size beyond 1000 neuron does not significantly change the oscillatory dynamics and dramatically increase computational costs. Pairs of interacting populations are connected in a sparse manner, following the method developed in Golomb and Hansel (2000): each neuron in a postsynaptic population *β* receives a number of synapses *K^sim^* from presynaptic population *α*, following this equation:

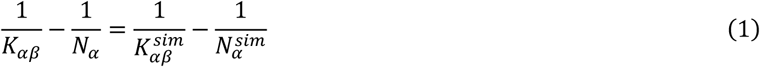

where *N_α_* is the size of population α (Table S1) and *K_αβ_* is the number of connections from population α to a neuron in population *β* (Table S2). The resulting *K_sim_* values used are shown in Table 1.

**Table 1.** Number of presynaptic neurons converging onto a postsynaptic neuron 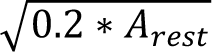. The values were computed from the population sizes and connectivity rates reported in the Supplemental Material (Tables S1, S2). Values that could not be estimated and were generalized from the other species model are marked by an asterisk.

| Presynaptic population | Postsynaptic population | Rat model | Monkey model |
| --- | --- | --- | --- |
| Arky | D1 | 28 | 28* |
|  | D2 | 28 | 28* |
| Cortex | D1 | 909* | 909 |
|  | D2 | 909* | 909 |
|  | FSI | 909* | 909 |
|  | STN | 52 | 92 |
| D1 | GPI | 980* | 980 |
| D2 | Proto | 757 | 954 |
| FSI | D1 | 79 | 79* |
|  | D2 | 53 | 53* |
| GPI | Thalamus | 333* | 333 |
| Proto | Arky | 94 | 94* |
|  | FSI | 32 | 32* |
|  | GPI | 234 | 234* |
|  | Proto | 94 | 94* |
|  | STN | 309 | 309* |
| STN | GPI | 446* | 446 |
|  | Proto | 60 | 603 |
| Thalamus | Cortex | 500* | 500 |

### Rate model

#### Dynamics

Neuron dynamics are described by a rate model (Wilson and Cowan, 1972), which allows to analytically investigate the dynamics of the different loops. Each neuron *i* is represented by its instantaneous activity which is given by *A_i_*(*t*) = *F*(*Isyn_i_*(*t*) + *Iext_i_* + *η*(*t*) − *θ*) where *F* is a piece-wise linear function with *F(x) = x* for *x* > 0 and 0 otherwise, and *θ* is a fixed activation threshold value of 0.1. The baseline drive *Iext_i_* is set such that *A_i_(t)* is equal to the instantaneous activity at rest *A*_rest_ (Table 2) under constant synaptic input *Isyn_i_*. The synaptic noise *η(t)* is modeled as a time-correlated, Ornstein–Uhlenbeck process (Gillespie, 1996):

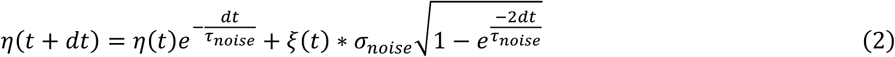

**Table 2.** Populations instantaneous activities at rest *A*_rest_ (mean ± s.d.). The mean firing rate of neurons in the various populations of the model where derived from estimates available in the experimental literature.

| Population | Values in the rat model (Hz) | Reference | Values in the monkey model (Hz) | Reference |
| --- | --- | --- | --- | --- |
| Arky | $14.1 \pm 8.3$ | (De la Crompe et al., 2020) | $14.21 \pm 1.04$ | (Katabi et al., 2023) |
| Cortex | $2.8 \pm 0.6$ | (De la Crompe et al., 2020) | $11.6 \pm 9.9$ | (Pasquereau and Turner, 2011) |
| D1 | $1.05 \pm 0.5$ | (Sharott et al., 2017) | $1.14 \pm 0.76$ | (Bogenpohl et al., 2013) |
| D2 | $0.49 \pm 0.4$ | (Sharott et al., 2017) | $1.14 \pm 0.76$ | (Bogenpohl et al., 2013) |
| FSI | $3.7 \pm 1.1$ | (Mallet et al., 2005) | $6.21 \pm 3.96$ | (Bogenpohl et al., 2013) |
| GPI | $26.0 \pm 3.1$ | (Benhamou and Cohen, 2014) | $67 \pm 32$ | (Galvan et al., 2011) |
| Proto | $39.8 \pm 13.7$ | (De la Crompe et al., 2020) | $65.1 \pm 34.1$ | (Galvan et al., 2011) |
| STN | $7.0 \pm 5.8$ | (De la Crompe et al., 2020) | $27.6 \pm 17.8$ | (Galvan et al., 2014) |
| Thalamus | $16.5 \pm 3$ | (Nakamura et al., 2021) | $18.2 \pm 10$ | (Anderson et al., 2003) |

where *τ_noise_* = 0.01s, *ξ*(t) is sampled from a standard normal distribution, and σ_noise_ is a noise amplitude relative to the neuron’s *A*_rest_ value, equal to 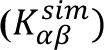. The sum of all synaptic input *Isyn^β^_j_* to a neuron *j* in population *β* is given by the following equation:

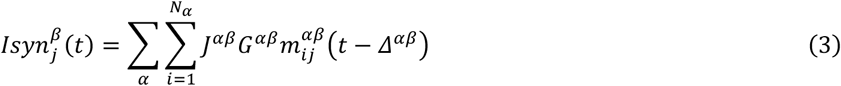

where *Δ^αβ^* is the synaptic delay from the presynaptic population *α* to population *β* (Table 3), *G^αβ^* is the synaptic weight, and J*^αβ^*is the connection matrix from population *α* to population *β*: this is a binary matrix of size 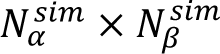 (in this case 1000×1000) whose entries are set to 1 (synapse present) with probability 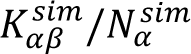 and to 0 otherwise, so that each neuron *j* receives on average 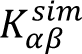 connections from population α. The synaptic contribution *m^α^* at each synapse is a low-pass filter of *A^α^* (Shriki et al., 2003) governed by the following equation:

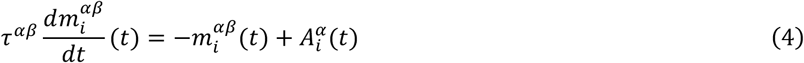

**Table 3.** Mean synaptic delays *Δ* used in the rate models (s.d. is set to 10% of the mean). The mean synaptic delays were derived from estimates in the experimental literature whenever available. For the rat model, studies performed in mice were also considered when rat data was not available. When no experimental estimate was available, the synaptic delay was set to its most likely value given the estimated delay in other species and the distance travelled by the axons: short axons for interactions within a nucleus, long axons between striatum or cortex and other BG nuclei, medium distance between non-striatal BG nuclei. NA: not available.

| Presynaptic population | Postsynaptic population | Mean delay, rat model (ms) | Reference | Mean delay, monkey model (ms) | Reference |
| --- | --- | --- | --- | --- | --- |
| Arky | D1 | 4.9 | (Glajch et al., 2016) | 10 | NA |
|  | D2 | 4.9 | (Glajch et al., 2016) | 10 | NA |
| Cortex | D1 | 12.8 | (Mallet et al., 2005) | 10.2 | (Nambu et al., 2002) |
|  | D2 | 12.3 | (Mallet et al., 2005) | 10.2 | (Nambu et al., 2002) |
|  | FSI | 8.3 | (Mallet et al., 2005) | 10.2 | (Nambu et al., 2002) |
|  | STN | 5.5 | (Degos et al., 2008) | 5.8 | (Nambu et al., 2000) |
| D1 | GPI | 7.2 | (Kita, 2001) | 13.1 | (Kita et al., 2006) |
| D2 | Proto | 6.89 | (Ketzel and Silberberg, 2021) | 10.5 | (Kita et al., 2006) |
| FSI | D1 | 0.84 | (Gittis et al., 2010) | 2 | NA |
|  | D2 | 0.93 | (Gittis et al., 2010) | 2 | NA |
| GPI | Thalamus | 5 | NA | 5 | NA |
| Proto | Arky | 4.55 | (Ketzel and Silberberg, 2021) | 5 | NA |
|  | FSI | 4.3 | (Glajch et al., 2016) | 10 | NA |
|  | GPI | 4.6 | (Ogura and Kita, 2002) | 4.6 | (Tachibana et al., 2008) |
|  | Proto | 4.7 | (Ketzel and Silberberg, 2021) | 5 | NA |
|  | STN | 1.3 | (Kita et al., 1983) | 2.0 | (Polyakova et al., 2020) |
| STN | GPI | 1.7 | (Nakanishi et al., 1991) | 4.7 | (Nambu et al., 2000) |
|  | Proto | 2.8 | (Kita and Kitai, 1991) | 5.5 | (Nambu et al., 2000) |
| Thalamus | Cortex | 5 | NA | 5 | NA |

where *τ^αβ^* is the synaptic time constant of the synapses from cells of population *α* to cells of population *β*, which is adapted from values of synaptic decay time constants found in the literature (Table 4). Lastly, the dynamics are simulated with a time step dt = 0.1 ms.

**Table 4.** Synaptic time constants *τ* used in the rate models. The synaptic time constants were derived from estimates available in the experimental literature whenever available. For the rat model, studies performed in mice were also considered when rat data was not available. When no experimental estimate was available, the synaptic time constant was set to 5 ms as most fast inhibitory and excitatory transmission have a decay time constant close to 5 ms. Since no estimate is available for NHP, we set all synaptic time constants to 5 ms in the monkey model. NA: not available.

| Presynaptic population | Postsynaptic population | Time constant, rat model (ms) | Reference |
| --- | --- | --- | --- |
| Arky | D1 | 28 | (Baufreton, unpublished) |
|  | D2 | 28 | (Baufreton, unpublished) |
| Cortex | D1 | 5 | NA |
|  | D2 | 5 | NA |
|  | FSI | 5 | NA |
|  | STN | 4.48 | (Karube et al., 2019) |
| D1 | GPI | 5.2 | (Connelly et al., 2010) |
| D2 | Proto | 6.13 | (Sims et al., 2008) |
| FSI | D1 | 11.4 | (Koos et al., 2004) |
|  | D2 | 11.4 | (Koos et al., 2004) |
| GPI | Thalamus | 7.8 | NA |
| Proto | Arky | 4.91 | (Sims et al., 2008) |
|  | FSI | 7.8 | NA |
|  | GPI | 2.1 | (Connelly et al., 2010) |
|  | Proto | 10 | NA |
|  | STN | 7.8 | (Baufreton et al., 2009) |
| STN | GPI | 1.8 | NA |
|  | Proto | 1.8 | Baufreton (unpublished) |
| Thalamus | Cortex | 5 | NA |

#### Parameters

The models were constrained by electrophysiological data extracted from the literature. For the rat model, most values were taken from a previous model by Azizpour Lindi, et al. (2024). We prioritized data from in vivo studies, and we aimed to use several values from the same study whenever possible to limit the imprecision due to different experimental methods. The instantaneous activity at rest *A_rest_* is normally distributed (mean and s.d. values in Table 2) and truncated at 0. The synaptic delays (Table 3) are normally distributed, using s.d. values equal to 10% of the mean. The synaptic time constants (Table 4) are adapted from synaptic decay time constants and are not distributed. Some parameters used in the rat model are taken from mouse studies when the data was not available in rats, they are indicated with “(mouse)”. Values unavailable in the NHP literature were set arbitrarily based on other rat and monkey parameters.

#### Theoretical analysis

The dynamics described previously allow to analytically derive the minimal conditions for the emergence of sustained stable oscillations in an isolated negative feedback loop of our model, as well as their frequencies at the bifurcation point. The details for the derivations are described in (Azizpour Lindi et al., 2024). The system exists either in a stable oscillatory state, in which oscillations do not decay over time, or a steady state, in which any oscillation caused by a perturbation eventually decays. The transition between these two states is determined by the absolute value of the product of the synaptic weights involved in the loop: we refer to this product as *G\*, and to the limit value separating the two states as *Glim*. The system is in the steady state when |*G* ∗ | < *Glim*, and in the stable oscillatory state when |*G* ∗ | > *Glim*. We expose below the derived formulas for the computations of *Glim* as a function of the loop synaptic delays Δ and time constants τ.

For a feedback loop with a single node α, such as the Proto loop:

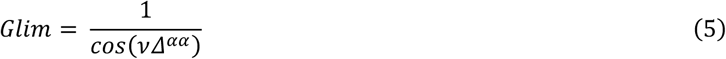

such that

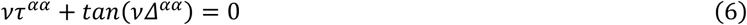

For a feedback loop with two nodes α and β, such as the STN loop:

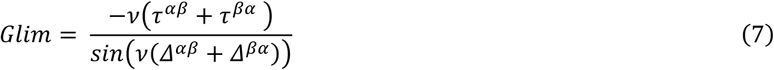

such that

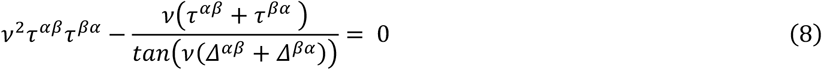

For a feedback loop with three inhibitory nodes α, β and γ, such as the FSI loop and the Arky loop:

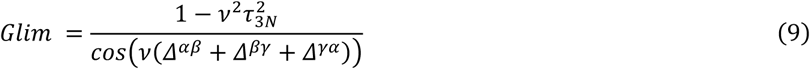

such that

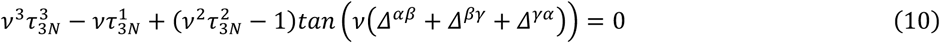

where

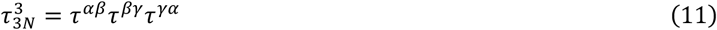

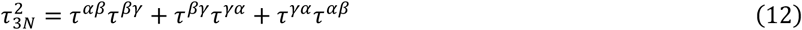

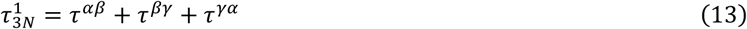

Lastly, for a feedback loop with four nodes α, β, γ and δ, such as the hyperdirect loop:

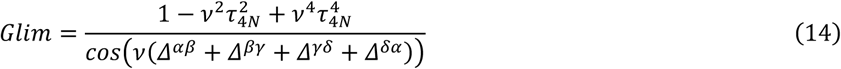

such that

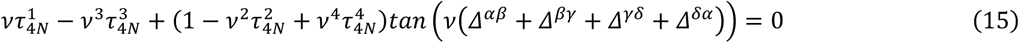

where

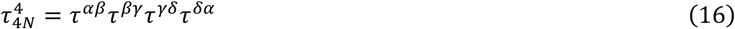

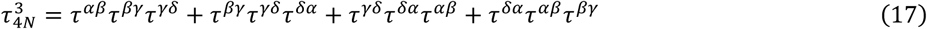

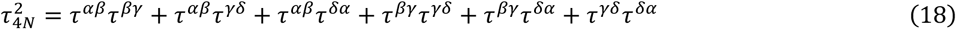

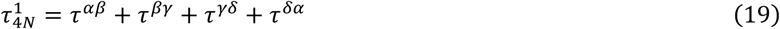

For all these cases, the resulting theoretical frequency of oscillations for |*G* ∗ | = *Glim* is equal to 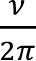. The polynomial root *ν* of each of the equations above was solved using the *scipy.optimize.fsolve* function in Python.

#### Simulation details

To reproduce the cortical stimulation protocol described in Kita and Kita (2011) and Nambu et al. (2000), an electrical cortical stimulation was simulated by artificially increasing the *Iext* of all Ctx neurons to 1500 during 0.3 ms. The stimulation was repeated 5 times, with an inter-stimulus interval of 1.0 s with the rat model and 1.5 s with the monkey model. We then processed the individual activities of the neurons to extract the latencies of onsets of excitation and inhibition events, taking the 200 ms before the stimulation as the baseline activities. As done by Kita and Kita (2011), the event onsets were taken as the first of 3 consecutive PSTH bins (bin size = 1 ms) above (early and late excitations) or below (short inhibition) the 95% confidence interval of the baseline. Results from the rat model were compared with those published by Kita and Kita (2011), and results from the monkey model were compared with those described by Nambu et al. (2000). Based on the populations observed in the two studies, the nuclei of interest were the Proto (to compare with GPe experimental response), the STN, and the GPi. We also grouped the onsets of the D1 and D2 of our rat model, to compare with the striatal response described by Kita and Kita (2011).

To simulate a pharmacological blockade of the STN, we performed a strong negative stimulation of all the STN cells, setting their *Iext* to -1000 over the entire duration of the simulation. For the pharmacological blockade of the GPe, this same forced inhibition was applied to 70% of the Proto and 70% of the Arky neurons, leaving the *Iext* of the remaining 30% unaltered, to reproduce the extensive but incomplete pallidal inactivation typically observed experimentally (Soares et al., 2004).

To selectively place one loop at a time in the oscillating state, the synaptic weights of that loop were set to *Glim^1/N^ + 0.3* where N is the number of nodes in the loop, while all other synaptic weights involved in the remaining generators were set to 0.2. The connections not involved in any generator were set to 1 (or -1). We simulated 4.5 seconds of activity, discarding the first 500 ms.

To analyze the effect on oscillations of the interaction between several feedback loops, we set all synaptic weights in the loops of interest to 1 (or -1) and varied the weight of one projection per loop: G_Proto–Proto_, G_STN–Proto_, G_D2–Proto_, and G_GPi–Th_. To clearly distinguish the steady from the oscillatory states, the synaptic noise was set to zero. The population sizes were lowered to 100 neurons per population to reduce the duration of the simulations. We simulated 5 seconds of activity and discarded the first 2 seconds to account for the adaptation period.

### Spiking neural network

A leaky integrate-and-fire (LIF) model of the rat BGTC network was implemented based on the work by Azizpour Lindi et al. (2024), using the same architecture (including population sizes *N^sim^* and connection probabilities *K^sim^*, see Eq. 1), average population activities (Table 2) and time step dt = 0.1 ms as the rat rate model.

#### Dynamics

The membrane potential *V* of each neuron is given by

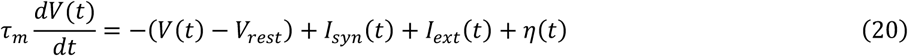

where *V_rest_* and *τ_m_* are respectively the resting membrane potential and the membrane time constant (Table 5). *I_syn_* is the total synaptic input, *I_ext_* is the external input to the neuron and *η* is the synaptic noise. The membrane potential is integrated at each time step using the second-order Runge–Kutta (RK2) method. When the membrane potential reaches the firing threshold *V_th_*, a spike is recorded and *V(t)* is set close the neuron’s *V_rest_* value (Table 5), using a linear interpolation method (Hansel et al., 1998) such that

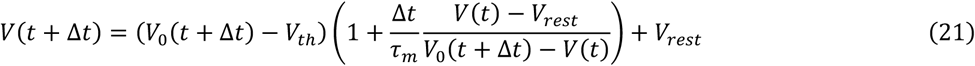

**Table 5:** Neuronal parameters used in the LIF model. Values are derived from the experimental literature and are given as mean ± s.d [min, max]). When two references are given, the first was used for the resting and threshold potentials, and the second for the membrane time constant.

| Population | $V_{rest}$ (mV) | $V_{th}$ (mV) | $\tau_m$ (ms) | References |
| --- | --- | --- | --- | --- |
| Arky | $-70 \pm 1$ [-90, -60] | $-55 \pm 2$ | $19.9 \pm 3$ [2, 100] | (Cooper and Stanford, 2000; Abdi et al., 2015) |
| Ctx | $-73.9 \pm 1.5$ [-90, -55] | $-42.8 \pm 1.63$ | $21.5 \pm 1.38$ [10,30] | (Perez-García et al., 2021) |
| D1 | $-76.8 \pm 3$ [-100, -55] | $-50 \pm 0.6$ | $4.9 \pm 0.5$ [2, 12] | (Slaght et al., 2004) |
| D2 | $-76.8 \pm 3$ [-100, -55] | $-50 \pm 0.6$ | $4.9 \pm 0.5$ [2, 12] | (Slaght et al., 2004) |
| FSI | $-78.2 \pm 0.5$ [-85, -60] | $-52.4 \pm 0.5$ | $3.1 \pm 0.3$ [1, 6] | (Schulz et al., 2011) |
| GPI | $-52.7 \pm 2$ [-90, -45] | $-38.8 \pm 1.6$ | $7.3 \pm 1.06$ [2, 15] | (Oh et al., 2017; Shin et al., 2017) |
| Proto | $-65 \pm 1$ [-85, -60] | $-54.8 \pm 1$ | $12.9 \pm 1.3$ [2.25, 45] | (Abdi et al., 2015; Karube et al., 2019) |
| STN | $-59 \pm 0.5$ [-75, -55] | $-50.8 \pm 0.5$ | $5.1 \pm 0.6$ [2, 10] | (Paz et al., 2005) |
| Th | $-60.4 \pm 1.35$ [-70, -52] | $-42.4 \pm 0.6$ | $12.2 \pm 1.1$ [6, 25] | (Paz et al., 2007; Jhangiani-Jashanmal et al., 2016) |

where *V_0_*(*t*+Δ*t*) is given by the RK2 at the spike time.

The external input *I_ext_* is set to reproduce the desired population firing rate (Table 2), interpolated from the population I–F curve (input–firing rate) computed beforehand. The synaptic noise is implemented with the same Ornstein–Uhlenbeck process and amplitude as in the rate model (Eq. 2).

Synaptic inputs are modeled by a double exponential with independent rise and a decay time constants (Fourcaud and Brunel, 2002). The sum of synaptic inputs *I_syn_*(*t*) from all presynaptic neurons of population *α* to each neuron *j* of population *β* is given by

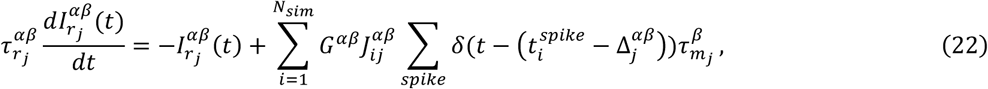

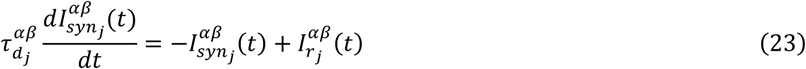

where 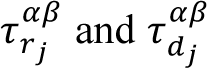 are the rise and decay time constants of the synaptic input from population *α* to neuron *j*, and 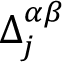 is the synaptic delay from population *α* to neuron *j* (Table 6). The connectivity matrix *J^αβ^* uses the same implementation as the rate model. The slope of the I–F curve is roughly 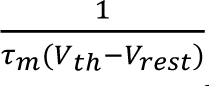 its linear section, which allows to approximate the synaptic weight *G^αβ^* from the rate model weight such that 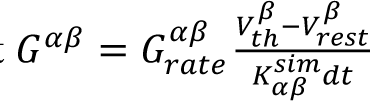 to reproduce the effective input-output gain.

**Table 6:** Synaptic parameters used in the LIF model. Values are derived from the experimental literature when available and given as mean ± s.d [min, max]). Estimates relied on studies performed in mice when rat data was not available. NA: no experimental estimate available.

| Presynaptic population | Postsynaptic population | $\tau_r$ (ms) | $\tau_d$ (ms) | Reference | $\Delta$ (ms) | Reference |
| --- | --- | --- | --- | --- | --- | --- |
| Arky | D1 | $1 \pm 0.5$<br>[0.1, 3] | $28 \pm 5$<br>[0.5, 35] | Baufreton (unpublished) | $4.9 \pm 0.6$<br>[3.8, 7] | (Glajch et al., 2016) |
| | D2 | $1 \pm 0.5$<br>[0.1, 3] | $28 \pm 5$<br>[0.5, 35] | Baufreton (unpublished) | $4.9 \pm 0.6$<br>[3.8, 7] | (Glajch et al., 2016) |
| Cortex | D1 | $1 \pm 0.5$<br>[0.1, 3] | $5 \pm 2$<br>[1, 10] | NA | $12.8 \pm 0.5$<br>[1.8, 10] | (Mallet et al., 2005) |
| | D2 | $1 \pm 0.5$<br>[0.1, 3] | $5 \pm 2$<br>[1, 10] | NA | $12.3 \pm 0.5$<br>[1.8, 10] | (Mallet et al., 2005) |
| | FSI | $1 \pm 0.5$<br>[0.1, 3] | $6 \pm 2$<br>[1, 10] | NA | $8.3 \pm 0.5$<br>[1.8, 10] | (Mallet et al., 2005) |
| | STN | $0.75 \pm 0.41$<br>[0.31, 0.77] | $4.48 \pm 2.12$<br>[1.26, 7.71] | (Karube et al., 2019) | $5.5 \pm 0.6$<br>[1, 10] | (Degos et al., 2008) |
| D1 | GPI | $1.2 \pm 0.1$<br>[0.2, 10] | $5.2 \pm 0.4$<br>[0.5, 30] | (Connelly et al., 2010)<br>(mouse) | $7.2 \pm 2.7$<br>[4.3, 11.3] | (Kita, 2001) |
| D2 | Proto | $0.8 \pm 0.22$<br>[0.2, 10] | $6.13 \pm 1.42$<br>[0.5, 30] | (Sims et al., 2008) | $6.89 \pm 0.35$<br>[4.3, 11.3] | (Ketzel and Silberberg, 2021) |
| FSI | D1 | $1.5 \pm 2.9$<br>[0.6, 3] | $11.4 \pm 2.1$<br>[0.5, 30] | (Koos et al., 2004) | $0.84 \pm 0.17$<br>[0.6, 2] | (Gittis et al., 2010) |
| | D2 | $1.5 \pm 2.9$<br>[0.6, 3] | $11.4 \pm 2.1$<br>[0.5, 30] | (Koos et al., 2004) | $0.84 \pm 0.17$<br>[0.6, 2] | (Gittis et al., 2010) |
| GPI | Thalamus | $1 \pm 0.2$<br>[0.1, 2] | $7.8 \pm 4.4$<br>[4.6, 18.4] | NA | $5 \pm 0.1$<br>[1.5, 10] | NA |
| Proto | Arky | $0.5 \pm 0.15$<br>[0.1, 10] | $4.91 \pm 1.08$<br>[0.1, 30] | (Sims et al., 2008) | $4.55 \pm 0.54$<br>[2.55, 7.05] | (Ketzel and Silberberg, 2021) |
| | FSI | $1.1 \pm 0.4$<br>[0.2, 10] | $7.8 \pm 4.4$<br>[0.5, 30] | Based on similarity to Proto–STN | $4.3 \pm 0.7$<br>[3.2, 7] | (Glajch et al., 2016) |
| | GPI | $0.41 \pm 0.02$<br>[0.1, 5] | $2.1 \pm 0.1$<br>[0.1, 5] | (Connelly et al., 2010) | $4.6 \pm 0.6$<br>[0.5, 8] | (Ogura and Kita, 2002) |
| | Proto | $0.5 \pm 0.15$<br>[0.1, 10] | $4.91 \pm 1.08$<br>[0.43, 6.86] | (Sims et al., 2008) | $4.7 \pm 0.88$<br>[3.05, 7.55] | (Ketzel and Silberberg, 2021) |
| | STN | $1.1 \pm 0.4$<br>[0.8, 1.6] | $7.8 \pm 4.4$<br>[4.6, 18.4] | (Baufreton et al., 2009) | $1.3 \pm 0.3$<br>[0.8, 2.5] | (Kita et al., 1983) |
| STN | GPI | $1 \pm 0.1$<br>[0.2, 10] | $1.8 \pm 2.5$<br>[0.43, 6.86] | Based on similarity to STN–Proto | $1.7 \pm 0.5$<br>[0.5, 5] | (Nakanishi et al., 1991) |
| | Proto | $0.6 \pm 0.1$<br>[0.2, 10] | $1.8 \pm 2.5$<br>[0.43, 6.86] | Baufreton (unpublished) | $2.8 \pm 0.6$<br>[2, 4.4] | (Kita and Kitai, 1991) |
| Thalamus | Cortex | $1 \pm 0.2$<br>[0.1, 2] | $5 \pm 1$<br>[2, 10] | NA | $5 \pm 1$<br>[2, 10] | NA |

#### Parameters used

Most values were taken from the reduced spiking model developed by Azizpour Lindi, et al. (2024), with additional parameters extracted from the literature. The population resting activities are the same as with the rat rate model (Table 2). All parameters are normally distributed across neurons from the same population, using a truncated normal distribution when a range is provided.

#### Simulation details

The generators were individually selected using the same approach as the rate models, by setting the synaptic weights of the loop of interest to high values and all other synaptic weights to low values. Spike times were extracted from 3.1-second simulations, and the first 100 ms were discarded to account for the initial adaptation period.

### Data analysis

#### Power spectrum

The power spectrum of the population average activity was derived with the *scipy.fft.fft* function, using a window size of 1 second, and normalized by the area under the curve to improve visualization.

#### Phase angle distribution

The phase histogram was calculated for a subset of five populations (Proto, Arky, D2, STN, and Ctx), using the Proto population as the reference for aligning the phases. The method differed between the rate and LIF models.

From the rate models simulations, we first averaged the coherence between 2000 random pairs of Proto neurons obtained with the *scipy.signal.coherence* function. The frequency between 1 and 100 Hz with the highest coherence magnitude was selected as the reference frequency. For each of the five populations of interest, we then derived the cross-spectral density (CSD) for 2000 random pairs of neurons between the selected population and the Proto population, which is the reference population, by applying the *scipy.signal.csd* function. For each pair of neurons, the imaginary part of the CSD value at the reference frequency was extracted as the phase angle.

The method for the LIF model simulations was adapted from previous analyses of neuronal spiking data (De la Crompe et al., 2020). The 10–50 Hz filtered activity of the Proto population mean activity was used as the reference signal. The phase angle distribution of each neuron of a population of interest was obtained by measuring the phase of each spike within the cycle created by the two closest bounding peaks of the reference signal. Subjecting all neurons to the Rayleigh test, only those which distribution was significantly not uniform were considered for the population phase histogram. The population phase histogram is the sum of the distributions of the selected neurons, normalized by the population firing rate.

Finally, a polar representation was used to plot the distribution of all the phase angles, with a bin size of 1°. The means and standard deviations of the angle distributions were calculated using the *scipy.stats.circmean* and *scipy.stats.circstd* functions respectively. For the rate model results, we used a logarithmic radial axis to improve visualization.

#### Oscillations from multiple feedback loops

To evaluate the stability of the oscillations in the population of interest (Proto or STN), we extracted the peaks of the average population activity from the last 3 seconds of the 5-s long simulations with the *scipy.signal.find_peaks* function in Python. If the signal had more than 10 peaks, with a prominence greater than 1.0, and if the amplitude of the highest peak in the second half of the signal was between 95% and 110% of the amplitude of the highest peak in the first half of the signal, we assumed the oscillations were stable. In that case, we extracted the main frequency using the *scipy.signal.welch* function.

### Code Accessibility

The custom Python code implemented for the model simulations and the data analyses is available on GitHub (https://github.com/ElenaNicollin/BG_rate_models)

## Results

### The rat and monkey BGTC network models

The properties of abnormal beta oscillations recorded in parkinsonian patients and in different animal models of the disease differ markedly. These properties have been extensively studied in both rodent (rat and mouse) and NHP models of PD. The anatomy and physiology of the BG network, well characterized in these species, are relatively consistent across mammals: anatomical organization is well conserved (Grillner and Robertson, 2016; Boraud et al., 2018; Kim, 2025), yet differences in brain size give rise to substantial differences in transmission delays, which may in turn affect the oscillatory properties of the network (Buzsáki et al., 2013). Here, we develop two models representing the dorsal part of the mammalian BGTC loop linking primary motor cortex to its BG targets, one for rats and one for monkeys. The two models share the same architecture derived from anatomical evidence (Alexander et al., 1986; Foster et al., 2021), focusing on the neuronal populations and circuits potentially involved in the generation of abnormal beta oscillatory activity in PD patients and animal models (Fig 1). Each model circuit comprises nine neuronal populations spanning the striatum, STN, globus pallidus, thalamus, and motor cortex. In the striatum, three inhibitory subpopulations represent, respectively: neurons expressing D1-like receptors and projecting mainly to BG output structures (GPi and SNr); neurons expressing D2-like receptors and projecting mainly to the GPe; and fast-spiking interneurons (FSIs). The globus pallidus is divided into external (GPe) and internal (GPi) segments in primates. In rodents, the globus pallidus as a whole is homologous to the primate GPe (Dudman and Gerfen, 2015); we therefore use GPe to refer to both primate GPe and rodent GP throughout. The GPe contains at least two distinct inhibitory neuronal populations, as evidenced by anatomical and physiological studies in rodents (Mallet et al., 2012), primates (DeLong, 1971; Sato et al., 2000; Katabi et al., 2023), and humans (Hutchison et al., 1994). Because these populations may play very different roles in PD oscillations (De la Crompe et al., 2020, 2025), they are represented separately in the model. One population, termed prototypic (Proto) neurons, sends projections to the STN and GPi; another, termed arkypallidal (Arky) neurons, projects exclusively to striatal projection neurons, as demonstrated in rodents (Mallet et al., 2012). Although the precise projection patterns of primate GPe neurons have not been fully characterized, some GPe neurons are known to project to the striatum (Sato et al., 2000), and indirect evidence links the two physiologically distinct populations of primate GPe neurons to Proto and Arky neurons (Katabi et al., 2023), suggesting conserved circuitry between rodents and NHPs. The primate GPi corresponds to the entopeduncular nucleus (EP) in rodents, whereas the substantia nigra pars reticulata (SNr) is a distinct output nucleus present in both species. In the model, a single inhibitory population, termed GPi, stands for the BG output nuclei as a whole (GPi and SNr in primates; EP and SNr in rodents). The STN, the ventral anterior and ventral lateral thalamic nuclei (Thalamus), and the primary motor cortex (Ctx) are each represented by a single excitatory neuronal population for simplicity. Within each population, neurons are modeled as rate units in which the dynamical variables describe synaptic activity (Wilson and Cowan, 1972; Shriki et al., 2003). The connectivity architecture reflects the principal known anatomical connections of the rodent and primate BG networks (Parent and Hazrati, 1995a, 1995b; Bolam et al., 2000; Foster et al., 2021).

Five negative feedback loops that may generate oscillatory activity emerge from the synaptic connections between these neuronal populations and are represented with different colors in Fig 1 (Pavlides et al., 2015; Azizpour Lindi et al., 2024; Zang et al., 2024). The Proto loop comprises a single node and represents lateral inhibition among Proto neurons. The STN loop corresponds to Proto inhibition onto the STN and its reciprocal excitatory projection back to the GPe. The inhibitory interactions among Proto, FSI, and D2 cells form the FSI loop, while those among Proto, Arky, and D2 cells form the Arky loop. Finally, the hyperdirect loop consists of cortical excitatory projections to the STN, which are relayed to the GPi, which in turn inhibits the thalamus, which projects back to the cortex.

For each negative feedback loop, neuronal dynamics depend strongly on the strength of the synaptic connections along the loop, with oscillatory activity emerging when feedback is sufficiently strong (see Materials and Methods for a detailed theoretical analysis). Because coupling strength along a given neuronal pathway is difficult to extract from experimental data, synaptic weights across all connections are varied throughout the simulations described below. In particular, we investigate conditions under which one or several loops operate in the oscillatory regime. Once in this regime, the properties of the resulting oscillatory dynamics depend critically on the temporal parameters of the circuit, namely the synaptic delays (*Δ_α_*, *_β_*) and synaptic decay times (*τ_α_*, *_β_*) between all connected population pairs {*α*, *β*} (see Materials and Methods). Wherever available, experimentally measured values of these parameters are implemented in both models. Together with population firing rate distributions, these values are drawn from electrophysiological data reported in the literature and are summarized in Tables 2, 3, and 4. Data from in vivo studies were prioritized and in vitro data were used when in vivo estimates were unavailable. Because the two models share the same architecture, they instantiate the same five negative feedback loops; however, their different temporal parameterizations shape the properties of the oscillatory activity that can emerge.

### Response of BG populations to cortical stimulation

While the literature offers a large body of data to constrain the models, it stems from heterogeneous experimental conditions (in vivo vs in vitro, rodent data mixing data from rats and mice) and yields an incomplete parameter set — particularly for NHPs, where the absence of in vitro recordings makes synaptic time constant data very scarce. We therefore sought to verify that the set of synaptic time constants and delays considered here accurately captures the temporal features of interactions between the various neuronal populations of the network. To this end, we compared the time course of responses in various BG neuronal populations following brief electrical stimulation of the motor cortex in the models with experimental reports of analogous responses in the literature. Responses of BG neuronal populations to cortical stimulation have been reported in both rats (Maurice et al., 1999; Kita and Kita, 2011) and monkeys (Nambu et al., 2000; Tachibana et al., 2008). Brief high-intensity cortical stimulation induced a multiphasic response in all populations of both models (Fig 2A). The average onsets of the early excitation, short inhibition, and late excitation in the GPe, STN, GPi, and striatum, obtained with the rat and monkey models, were compared directly to values reported in experimental studies (Fig 2B; Table S3). To disentangle the contributions of the different pathways to these temporal dynamics, we then simulated local pharmacological inactivation of the STN and GPe and examined their effects on the GPi response to cortical stimulation (Fig 2C), following experimental protocols reported in the literature (Maurice et al., 1999; Nambu et al., 2000; Tachibana et al., 2008).

**Figure 2.**
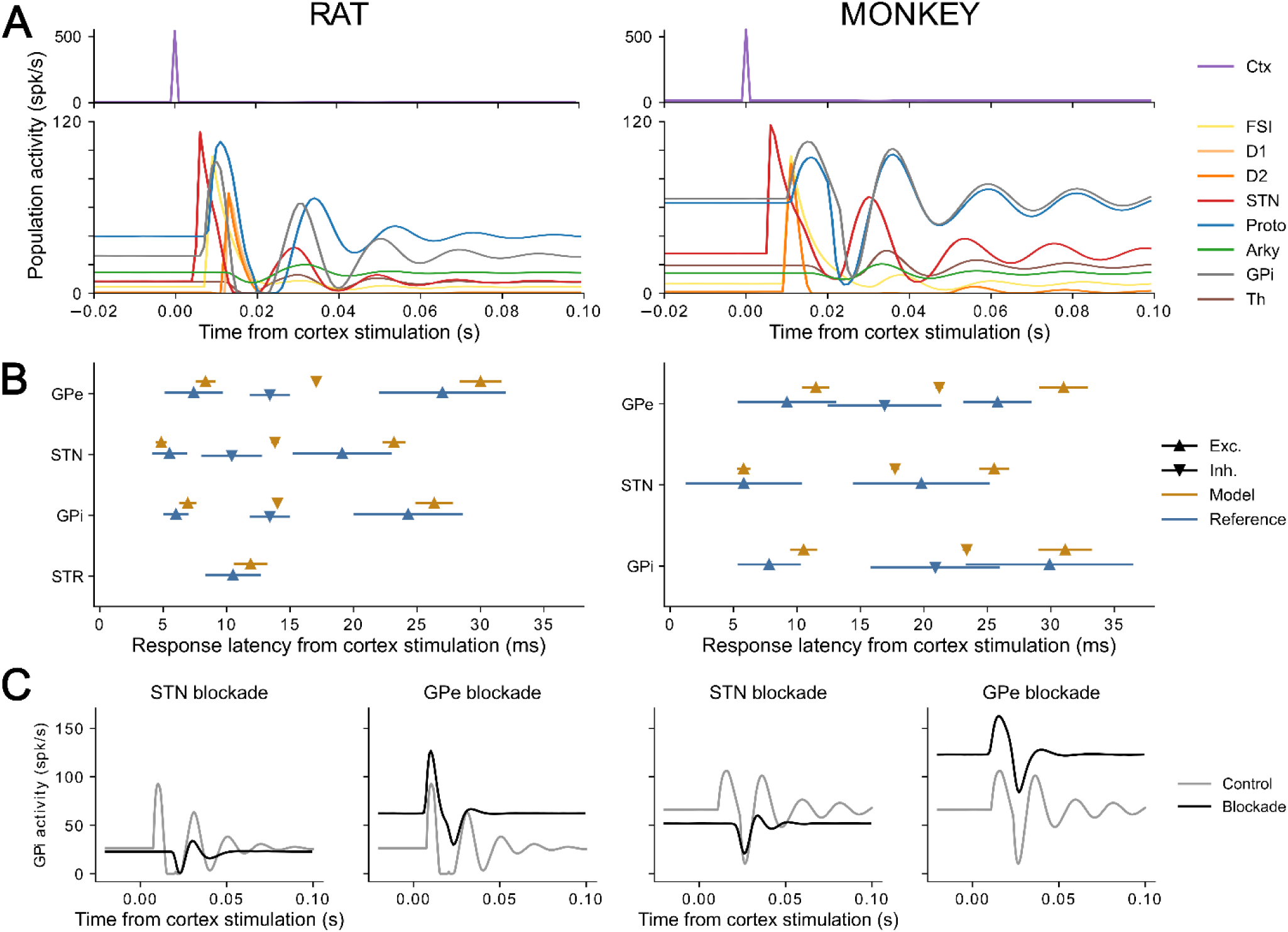
Temporal profile of the response to cortical stimulation in various populations of the BGTC model, compared to experimental data. A: Average activities of the different BGTC neuronal populations over n = 5 trials, centered on the time of cortical stimulation in the rat (left) or monkey (right) model, with a time bin of 1 ms. B: Mean (triangles) and standard deviation (lines) of the latencies of excitation (Exc.) and inhibition (Inh.) onsets in selected BG populations, obtained with the rat (left, yellow lines and triangles) and monkey (right, yellow lines and triangles) models and reported in reference experimental papers (blue lines and triangles). C: Effect of STN or GPe pharmacological blockade on GPi population response to the cortical stimulation. Gray line: response of GPi neurons before STN or GPe pharmacological blockade, black line: response after STN or GPe pharmacological blockade.

Cortical stimulation first drives short-latency excitation in the STN and striatum, both of which receive direct cortical input. The resulting increase in STN activity drives the initial rise in activity in the GPe and GPi, and STN inactivation abolishes the early GPi excitation, consistent with experimental findings (Maurice et al., 1999; Nambu et al., 2000; Tachibana et al., 2008). Rapid activation of GPe Proto neurons in turn induces a brief inhibition of STN neurons. The combination of increased inhibitory input from striatal neurons and reduced excitatory drive from the STN then produces a period of inhibition in both GPe and GPi. In the rat model, GPi inhibition is slightly delayed following either STN or GPe inactivation, suggesting that under normal conditions it is initially driven by the STN-to-GPe pathway and subsequently sustained by direct D1 striatal inhibition. In the monkey model, by contrast, neither inactivation affects the timing of GPi short inhibition, implying that D1-mediated inhibition arrives no later (and possibly earlier) than the inhibition relayed via the STN-to-GPe pathway. The transient inhibition of pallidal neurons ultimately leads to disinhibition of the STN by GPe Proto neurons, driving a late excitation in the STN which in turn produces late excitations in the GPe and GPi. This late GPi excitation is nearly abolished in both models when activity in the STN or GPe is suppressed.

The simulated event onset timings (see Methods for the calculation of onset latencies) are overall consistent with experimental observations, with a few exceptions. In the rat model, the STN short inhibition onset occurs at 13.7 ± 0.2 ms, whereas Kita and Kita (2011) reported a latency of 10.4 ± 2.3 ms. Reducing the pallido-subthalamic synaptic delay from 1.3 ms (as reported in (Kita et al., 1983)) to 0.5 ms did not close this gap (13.1 ± 0.3 ms, Fig S1). The shorter inhibition latency observed experimentally may reflect spike-frequency adaptation in STN neurons (Wilson et al., 2004; Barraza et al., 2009), which could accelerate the post-excitation decline in STN activity and thereby shorten the inhibition onset. Incorporating spike-frequency adaptation into STN neurons modestly reduced the inhibition latency (12.5 ± 0.4 ms; Fig S1) but also delayed the onsets of late excitation in the STN, GPe, and GPi. For these reasons, we chose to work with the non-adapting rate model which, while unable to fully capture some of the fast dynamics observed experimentally, reproduces longer-timescale effects most accurately, including the late excitation onsets. In the monkey model, event onsets in the GPe, STN, and GPi are slightly slower than, but broadly consistent with, those reported by Nambu et al. (2000). The GPe late excitation onset from the model (31.0 ± 1.9 ms) appears delayed relative to that reference (25.8 ± 2.6 ms), but falls within the range reported in other stud ies (30.8 ± 1.9 ms (Kita et al., 2004); 28.1 ± 4.4 ms (Iwamuro et al., 2017)). The STN neurons in our model showed a slight but significant short inhibition (17.8 ± 0.2 ms) which was not captured experimentally, although the authors do report a brief period of inhibition separating the STN early and late excitations. This inhibition is rarely significant in experimentally reported responses, likely due to the large fluctuation in baseline firing rate it is compared to. The variability of response latencies across neurons within a given population is lower in the model than observed experimentally, likely due to the simplified representation of neurons as firing rate units with rectified linear input-output function and limited sources of heterogeneity (distributed firing rates, synaptic delays, and amplitude of synaptic noise).

Overall, these results demonstrate that our rate models can reproduce the species-specific temporal signatures of cortical information transmission along the BGTC network, and confirm that both models provide an accurate representation of the network’s temporal properties.

### Theoretical constraints on the generated oscillations of each feedback loop

Oscillations can arise from the spontaneous activity of neuronal populations interconnected in a negative feedback loop (Ermentrout et al., 2001; Zang et al., 2024). Five such loops capable of generating oscillations are present in our BGTC models (Fig 1). Each loop is a dynamical system that, under constant external input, settles into either a steady state in which all populations display constant firing rates, or an oscillatory limit cycle in which the firing rates of all populations oscillate synchronously. The transition from a stable steady state to the oscillatory regime is known as a Hopf bifurcation (Strogatz, 2018). By isolating a given loop from the full network, we can analytically derive its dynamical state (steady state vs. oscillatory limit cycle) as a function of the model parameters — that is, determine the minimal conditions for the emergence of oscillatory activity as well as the corresponding oscillation frequency (see Materials and Methods, Theoretical Analysis, for detailed calculations). The parameters governing the transition between these two states are the number of nodes in the loop, the synaptic delays {*Δ^α,β^*}, the synaptic time constants {*τ^α,β^*}, and the synaptic weights {*G^α,β^*} for all connections between populations α and β in the loop. Of these, the first three are already fixed in our models, determined from the experimental literature and validated in the previous section, leaving the synaptic weights as the only free variables. Furthermore, all synaptic weights within a given loop contribute to the oscillatory instability through their product *G*=|Π G^α,β^|* (see Material and Methods, Theoretical analysis). Specifically, a loop remains in a stable steady state if and only if *G\** is below a critical threshold *Glim*, while oscillatory activity emerges when *G* > Glim* (Fig 3A). Because shorter delays increase the *Glim*, loops with fewer nodes and thereby shorter overall feedback delays, such as the Proto loop where only one synaptic delay is involved, have a higher *Glim* threshold than loops with more nodes, where several delays add up. Importantly, the relative values of *Glim* across the five loops differ between the rat and monkey models, reflecting differences in their synaptic parameters. Because the rat model features faster transmission with shorter synaptic delays, the transition to oscillations occurs systematically at higher gain products *Glim*. This implies that, for identical synaptic weights, the same loop could be in a steady state in one model yet in an oscillatory state in the other. Moreover, *Glim* values are more spread across loops in the rat model; this is most pronounced for the three longest loops (FSI, Arky, and hyperdirect) which range from 1.5 (hyperdirect loop) to 2.7 (Arky loop) in the rat model, whereas they are similar across all three loops in the monkey model (1.2 < *Glim* < 1.3).

**Figure 3.**
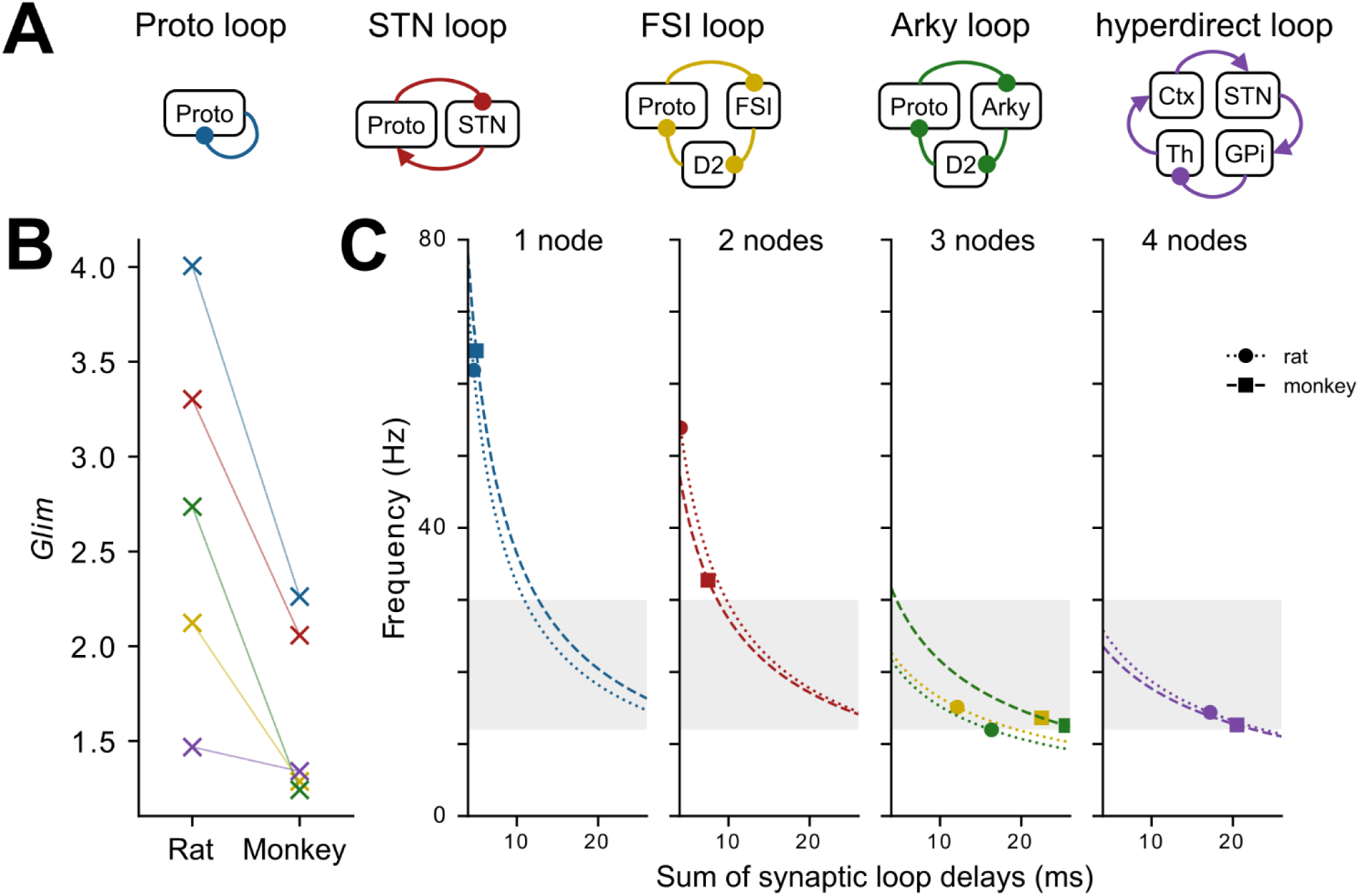
Theoretical derivations of limit gain and oscillation frequency in single oscillatory circuits. A: Negative feedback loops of interest isolated from the full network, color-coded for reference for all following figures. B: Critical product *Glim* of synaptic weights, delimiting steady and oscillatory states, derived for each individual negative feedback loop. The species-specific temporal properties of each loop network affect the value of *Glim*. C: Oscillation frequencies for negative feedback loops of different sizes, as a function of the sum of synaptic delays involved. Lines (i.e. dotted for rat, dashed for monkey) represent the predicted frequencies for each model for the full range of values of hypothetical synaptic delays; markers (i.e. round for rat, square for monkey) indicate the frequency predicted for the synaptic delays used in the models. The light grey band marks the beta range (13–30 Hz).

Beyond determining the critical threshold *Glim*, the synaptic time constants and delays along a given loop also impose the frequency of oscillations at the bifurcation (i.e., when *G\** = *Glim*), and this frequency does not vary much as *G\** further increases above that threshold, as shown in simulations (Fig 3). We computed the theoretical frequencies generated by each loop in both models as a function of the total sum of synaptic delays involved (Fig 3C), marking the values corresponding to the delays used in each model. It is worth noting that the scope of delays explored can extend beyond the realistic range, particularly for the shorter loops where only few short connections are involved. At the bifurcation, the two shortest generators, i.e. the Proto and STN loops, produce oscillation frequencies above the beta range: 62 Hz and 54 Hz, respectively, in the rat model, and 64 Hz and 32 Hz in the monkey model. By contrast, the three longer generators (Arky, FSI, and hyperdirect loops) oscillate in and below the low-beta range, between 11 and 15 Hz in both models. As noted above, these frequencies depend on the specific values of synaptic delays and time constants, which are derived from experimental data (when available). However, the theoretical analysis also reveals that as the number of nodes in a loop increases, the influence of the total synaptic delay on oscillation frequency decreases. In our BGTC network models, this implies that using high-confidence values for synaptic delays is most critical for the Proto loop (1 node) and the STN loop (2 nodes), and progressively less so for the FSI and Arky loops (3 nodes) and the hyperdirect loop (4 nodes), with respect to the frequency of the generated oscillations.

### Oscillation frequencies and phases from individual generators

The mathematical analysis of individual negative feedback loops provides, for each loop, the conditions on synaptic parameters for a stable steady state and the critical threshold for bifurcation into the oscillatory regime as detailed above. We next characterize the dynamics of the fully connected BGTC network in which all synaptic time constants and delays are fixed (Tables 3 and 4). Due to the dense interconnectivity of the network, interactions between loops give rise to network dynamics far more complex than the dynamics of isolated negative feedback loops. Interactions between coupled oscillators can produce a rich repertoire of network dynamics, including in-phase entrainment, cancellation of oscillations through phase opposition, emergence of new oscillation frequencies, and even chaotic behavior (Kuramoto, 1984). While the dynamics of the complete network cannot be fully determined analytically, its oscillatory properties can to a large extent be understood from the analysis of segregated loops conducted above. Before exploring the complex emergent dynamics arising from the interaction of several oscillating loop circuits, we first confirm that the oscillatory properties derived for isolated loops serve as a reliable proxy for oscillatory activity in the full network when at most one loop operates in the oscillatory regime (Fig 4). In particular, we verify that varying the synaptic weights along individual loops allows independent control of the dynamical state of each negative feedback loop. We begin from a configuration in which all synaptic gains are set to a low value of 0.2, such that each loop, if isolated, would remain in the steady-state regime (*G* < Glim*). In this configuration, all neurons display constant firing rates with no oscillatory fluctuations (Fig 4A, top left). Increasing the synaptic gains along a single loop well beyond its critical bifurcation threshold (*G\** > *Glim*) leads to the emergence of sustained synchronous oscillatory activity throughout the network (Fig 4A). Neuronal populations not belonging to the loop with increased gain also oscillate, driven by their synaptic connections from populations within that loop.

**Figure 4.**
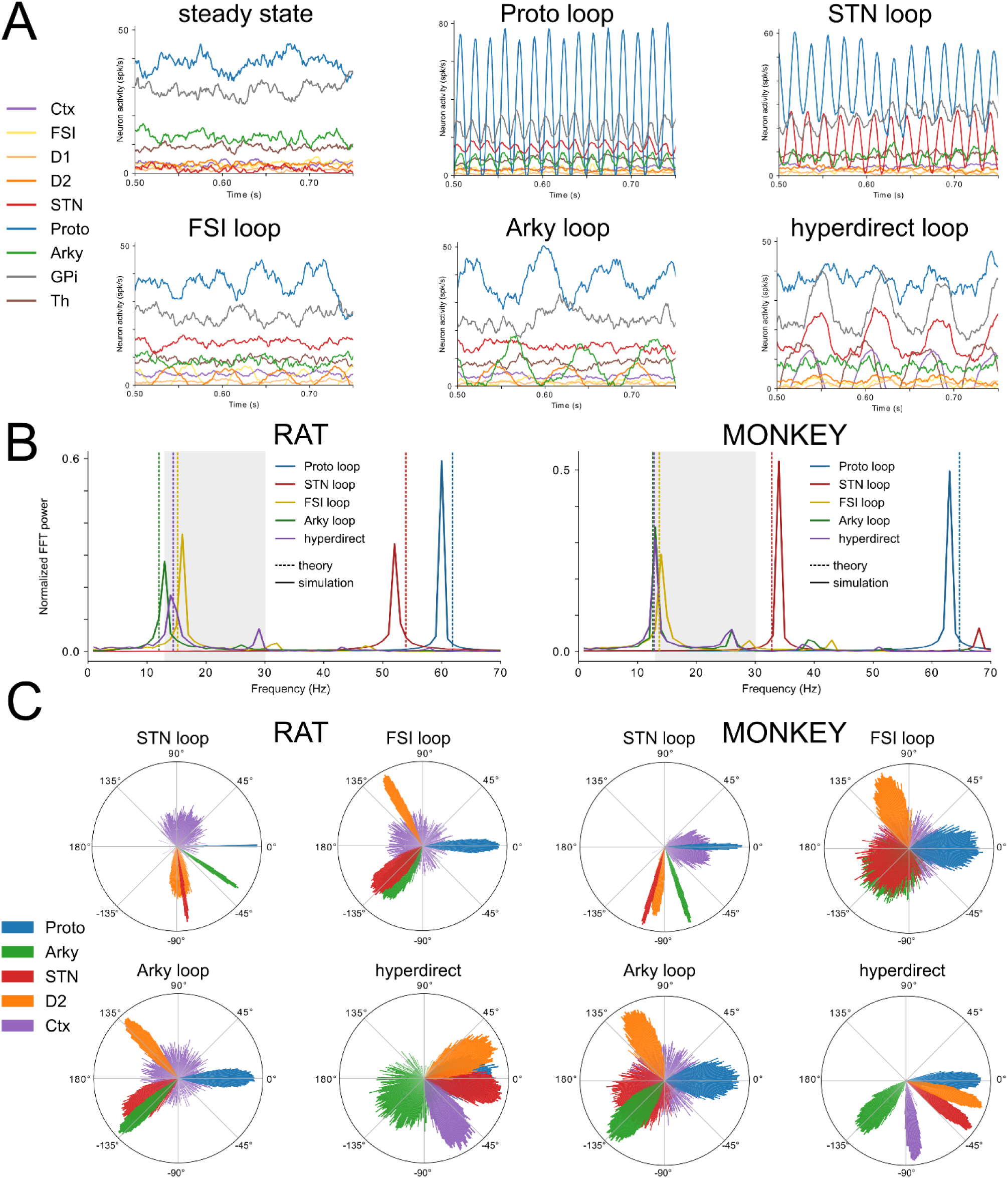
Power spectra and phase relationships generated by each negative feedback loop. A: Single neuron activities sampled from simulated BGTC network activities, with the network transitioning from steady, non-oscillatory state (top, left) to different oscillatory states as the synaptic weights of one negative feedback loop are increased. B: Power spectrum of the average activity in the Proto population from 4-s long simulations with the rat model (left) or the monkey model (right), with one negative feedback loop above the oscillatory threshold at a time (indicated as line color, see inset). Dashed lines indicate the theoretical frequencies derived previously. The light grey band marks the beta range (13–30 Hz). The Proto and STN loops generate a peak frequency above the beta range; the FSI, Arky and hyperdirect loops generate a peak frequency in the low-beta range. C: Polar histograms show the distribution of preferred firing phases of Proto, Arky, STN, D2 and Ctx neurons relative to Proto neurons (n = 2000 random pairs of neurons per distribution) from simulations in the rat (left) or monkey (right) model with specific negative feedback loops driving oscillations.

Using this approach, we characterized the frequency of oscillatory activity driven by each negative feedback loop separately within the full BGTC network, for both models (rat and monkey), through spectral analysis of the mean firing rate of the Proto population (Fig 4B). The peak frequencies obtained in these simulations are largely consistent with the theoretical predictions (Fig 3B). Three loops consistently generate oscillations in the low-beta range: the Arky loop (rat: 13 Hz; monkey: 13 Hz), the FSI loop (rat: 16 Hz; monkey: 14 Hz), and the hyperdirect loop (rat: 15 Hz; monkey: 13 Hz). The STN loop oscillates above the beta range, with a more pronounced difference between the two models (rat: 53 Hz; monkey: 33 Hz). The Proto loop oscillates at the highest frequencies (rat: 66 Hz; monkey: 62 Hz).

These results confirm that each loop can independently induce and sustain oscillations in the full network at frequencies close to those predicted analytically. Although several loops generate oscillations in a similar frequency range (e.g., the Arky, FSI, and hyperdirect loops all produce low-beta oscillations in both models), the exact activity patterns across the network appear to differ depending on which loop drives the oscillation (Fig 4A). To further discriminate the properties of oscillations generated by the different loops, we characterized the phase relationships between pairs of neurons drawn from distinct neuronal populations to capture the relative timing of their oscillatory coupling. In Fig 4C, we computed the phase distributions of neurons from the Cortex, STN, D2, Proto and Arky populations relative to Proto neurons, for oscillations driven by each negative feedback loop in both models. These phase distributions can be compared to phase differences measured experimentally from recordings between neuronal structures (Mallet et al., 2008a, 2012). Because different oscillation-generating loops produce distinct phase relationships between circuit components (De la Crompe et al., 2020), such differences provide an important signature for discriminating between candidate mechanisms underlying oscillation generation. We selected these populations from our models for their relevance in experimental works: these nuclei are commonly accessible in recordings of beta oscillations and typically display activities with strong beta synchronization from which well-defined phases can be extracted.

Figure 4C shows that phase angle distributions for a given loop are broadly similar between the rat and monkey models and differ more substantially depending on which loop generates the oscillatory activity. When the STN loop drives oscillations, the STN and Arky populations are tightly phase-locked (as seen from the narrow standard deviations) relative to the Proto population, with mean phase shifts of -82° ± 1° and -35° ± 2°, respectively, in the rat model, and -106° ± 1° and -72° ± 2° in the monkey model. The D2 population is phase-locked at -85° ± 10° and -99° ± 4° (rat and monkey model, respectively), and cortical neurons are phase-locked at 69° ± 41° and -6° ± 24° (rat and monkey model, respectively). With either of the two striatopallidal loops driving oscillations, phase relationships are broadly similar across both loops and both models, typically varying by 20° or less: D2 neurons are strongly phase-locked to Proto, with mean angles between 110° ± 10° (FSI loop, monkey model) and 131° ± 3° (Arky loop, rat model); STN neurons between -147° ± 40° (FSI loop, monkey model) and -139° ± 8° (FSI loop, rat model); and Arky neurons between -139° ± 3° (Arky loop, rat model) and -129° ± 38° (FSI loop, monkey model). Cortical neurons are not particularly phase-locked, with mean angles ranging from 77° to 144° relative to Proto and standard deviations of over 90°. Lastly, when the hyperdirect loop drives oscillations, the Proto, STN, and D2 populations oscillate nearly in phase: STN neurons show a mean angle of -10° ± 10° relative to Proto in the rat model and -36° ± 4° in the monkey model, while D2 neurons show mean angles of 27° ± 11° and -18° ± 4°, respectively, and Arky neurons are phase-locked at -133° ± 50° and -135° ± 9°, respectively. In contrast to the striatopallidal conditions, cortical neurons are strongly phase-locked to the Proto population (rat: -59° ± 10°; monkey: -83° ± 4°).

Taken together, these results highlight the discriminatory power of mean inter-population phase differences: loops that cannot be distinguished from their power spectrum alone, such as the hyperdirect loop and the two striatopallidal loops, can be clearly differentiated by the phase relationships among BG populations. Furthermore, neuronal populations that are part of the oscillation-generating loop tend to be more tightly phase-locked. This is illustrated by the striatopallidal results: while the mean phase angle of Arky neurons is similar across the Arky and FSI loop conditions, the phase distribution is narrower in the Arky loop condition, in which Arky neurons actively participate in oscillation generation, than in the FSI loop condition, in which their oscillatory activity merely reflects the oscillating output of the presynaptic Proto population.

### Conservation of oscillation properties in a spiking model of the rat BGTC network

Our firing rate model is a simplified representation of neuronal dynamics in the BGTC network, but the conservation of neuronal dynamics in higher complexity models is not always trivial (Devalle et al., 2017; Tse et al., 2026). We aimed to determine whether the oscillatory properties such as frequencies and phase shifts observed with that model are consistent with those produced by the same loop circuits in a spiking network model. To this end, we implemented a biologically constrained leaky integrate-and-fire (LIF) spiking model of the rat BGTC network, with cellular and synaptic parameters drawn from the literature (see Materials and Methods). While in vitro studies provide measurements of neuronal and synaptic properties in the rodent BGTC network, comparable data are largely unavailable for monkeys, restricting this implementation to the rat model exclusively.

Synaptic weights were initially set such that all neuronal populations were in an asynchronous state (Fig 5, top). As in the rate model, increasing synaptic weights along a negative feedback loop circuit (see Materials and Methods) led to the emergence of synchronized oscillatory activity across all populations belonging to that loop. We selectively placed each negative feedback loop in the oscillatory regime by adjusting the relevant synaptic weights and simulated the corresponding network activity (Fig 5). Oscillation frequencies were estimated from spectral analysis of the mean firing rate in each population (Fig 5A). The frequencies produced by the FSI loop (14 Hz), Arky loop (10 Hz), and hyperdirect loop (13 Hz) were consistent with those reported by the rate model. The STN loop, however, oscillated at 37 Hz in the spiking model, a substantial reduction from the 53 Hz observed with the rate model. This discrepancy likely reflects the slow membrane time constant of Proto neurons in the LIF model (12.9 ± 10.2 ms (Karube et al., 2019)), which is not captured by the single subthalamo-pallidal synaptic time constant used in the rate model (1.8 ± 2.5 ms), thereby slowing network dynamics. Despite this difference, the STN loop oscillation frequency remains largely above the beta range.

**Figure 5.**
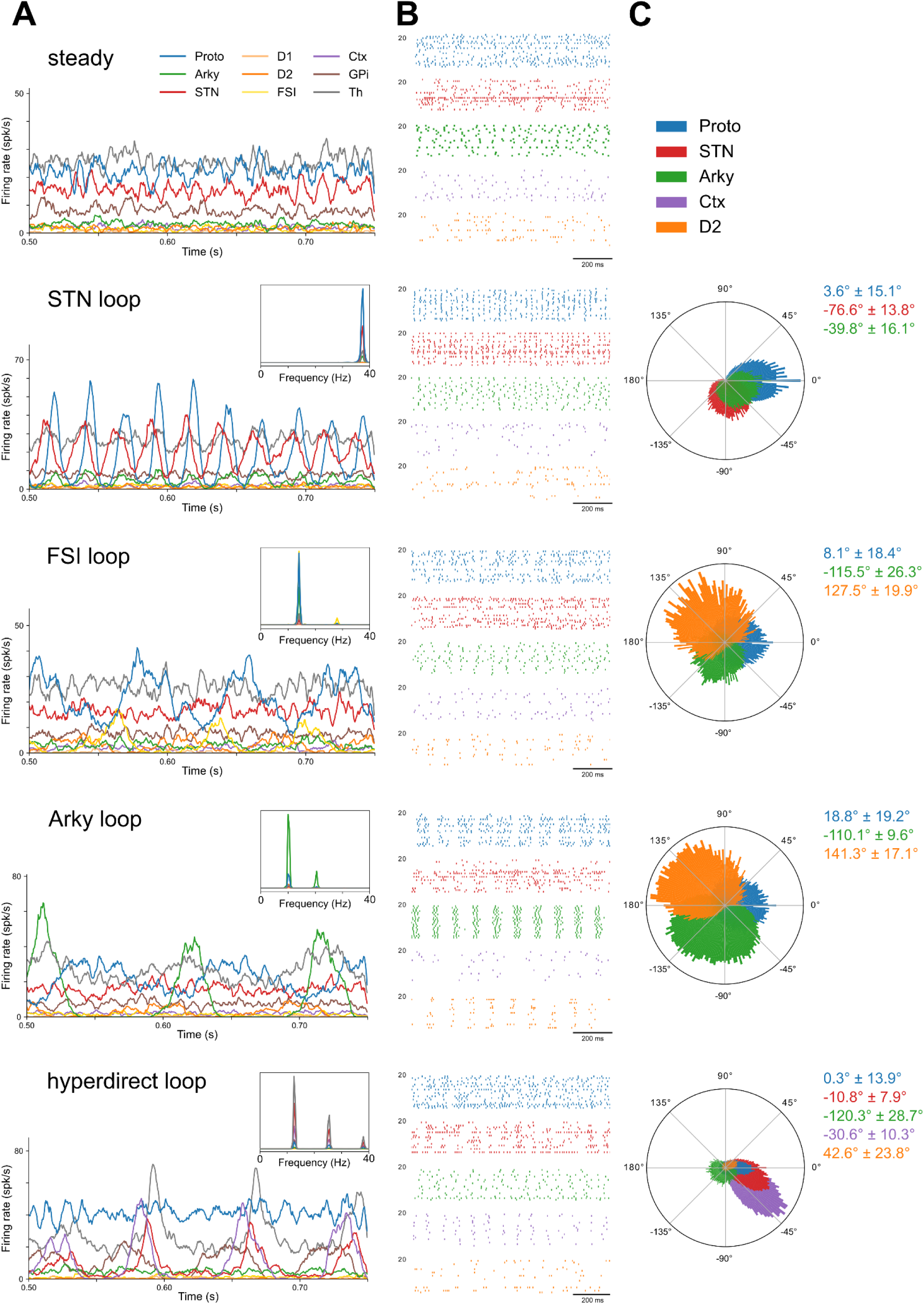
Neuronal dynamics and phase relationships generated by each negative feedback loop with the LIF spiking model of the rat BGTC network. A: Mean population activities of the rat BGTC network simulated with the LIF spiking model in the steady regime or with a negative feedback loop above the oscillatory threshold. Insets show the power spectra of the population activities over a 3-s long simulation for each driving generator. B: Raster plot of the spikes of n = 20 randomly chosen neurons from each of the Proto, STN, Arky, D2 and Ctx populations. Oscillatory synchronization appears between neurons within the populations as different generators drive oscillations. C: Polar distribution of the preferred firing phase among all Proto, STN, Arky, D2 and Ctx neurons relative to the Proto population activity. Only neurons that passed the Rayleigh test are included in the distributions, and the values of the mean and standard deviations are only reported for nuclei where over 20% of neurons passed the Rayleigh test.

Oscillatory synchronization was also apparent in the spiking activity of individual neurons within the different populations (Fig 5B). We computed the preferred firing phase of all neurons of the Proto, STN, Arky, D2 an Ctx populations relative to the reference signal established from the 10–50 Hz filtered Proto population activity (Fig 5C). The resulting phase shift distributions showed greater variability than in the rate model, likely owing to the multiple sources of heterogeneity in the LIF model (resting membrane potential, spike threshold, synaptic time constants, etc.). Nevertheless, the mean phase relationships between populations were remarkably consistent with those observed in the rate model. With the STN loop driving the oscillations, over 90% of neurons in the Proto, STN, and Arky populations were entrained. The relative phase angles among these populations were conserved compared to phases from the rate model, with STN neurons spiking on average at -76.6° ± 13.8°, followed closely by Arky neurons at -39.8° ± 16.1°. Meanwhile, less than 5% of D2 and Ctx neurons were significantly entrained. With either striatopallidal loop as the generator, the most strongly entrained populations were Proto, Arky, and D2 neurons, while only 3 to 7% of Ctx and STN neurons were significantly entrained. Entrained D2 neurons spiked at mean angles of 127.5° ± 19.9° (FSI loop, 38% of neurons) and 141.3° ± 17.1° (Arky loop, 59%), and Arky neurons at -115.5° ± 26.3° (FSI loop, 61%) and -110.1° ± 17.1° (Arky loop, 100%). With the hyperdirect loop as the generator, all five populations of interest were entrained (26% of D2, 68% of Arky, 79% of Proto, and 100% of STN and Ctx). The phase relationships were well conserved relative to the rate model: D2, cortical and STN neurons fired in-phase with Proto neurons, firing respectively at 42.6° ± 23.8°, -30.6° ± 10.3°, and -10.8° ± 7.9°, while Arky neurons were anti-phase to them, with a preferred firing phase of -120.3° ± 28.7°.

Taken together, the neuronal activities simulated with the spiking network model displays the same oscillatory properties as those obtained with the rate model. These results underscore the extent to which the intrinsic connectivity and temporal properties of a network — synaptic delays and time constants — jointly determine both the attainable oscillation frequencies and the phase angle distributions characterizing oscillations generated by specific negative feedback loops. Given the close agreement between the rate and spiking models of the rat BGCT network demonstrated here, we consider that oscillatory properties derived from the rate model can be meaningfully compared to experimental data, including in NHPs, for which the cellular and synaptic parameters required to build a biologically realistic spiking model are not yet available.

### Oscillations frequencies from interacting generators

Up to this point, the oscillatory properties of the network were characterized when a single loop circuit operated in the oscillatory regime. When two or more loops are simultaneously in the oscillatory regime, the resulting neuronal dynamics of the full BGTC network become more difficult to predict, as interactions between independent oscillators can produce a variety of behaviors, including sustained oscillatory activity, chaos, or stable asynchronous activity. To study how interactions between generators shape oscillation frequencies in the two animal models, we characterized the oscillatory properties of the rat and monkey BGTC rate models while varying four synaptic weights: {G_Proto–Proto_, G_D2–Proto_, G_STN–Proto_, G_GPi–Th_}. Each of these weights, when increased sufficiently, can switch a given loop from the asynchronous to the oscillatory regime: G_Proto–Proto_ controls the Proto loop, G_D2–Proto_ controls both striatopallidal loops, G_STN–Proto_ controls the STN loop, and G_GPi–Th_ controls the hyperdirect loop. The synaptic weights of all remaining connections in the network were set to 1. For each combination of synaptic weights, network activity was simulated for 3 s, and we report both the nature of the activity (asynchronous or oscillatory) and the peak oscillation frequency (when oscillatory; see Materials and Methods) in the Proto population (Figs. 6A and 7A) and in the STN population (Fig S2, Fig S3). The single-loop dynamics described above were recovered when all but one synaptic weight were set below their respective loops’ bifurcation thresholds. When two loops were able to drive oscillations simultaneously, the resulting network dynamics fell into one of three scenarios. In the first scenario, both loops contribute simultaneously to oscillation generation, and a shift in the relative feedback gain between the two loops produces a continuous shift in the dominant frequency. This is exemplified by the progressive increase in oscillation frequency driven by the striatopallidal loops as G_STN–Proto_ is increased and fast oscillations from the STN loop come into play, observed in both the rat (Fig 6Ba) and monkey (Fig 7Ba) models. Importantly, in this scenario, the pattern of phase-shifts between the various populations of the model changes only marginally, remaining most similar to the phases of the main driving oscillator. In the second scenario, the network undergoes a sharp transition from an oscillatory regime dominated by one loop to one dominated by the other as the relative loop strength is varied, resulting in a discontinuous jump in oscillation frequency. This is illustrated by the abrupt doubling of oscillation frequency from ∼13 to ∼26 Hz when oscillations coherently driven by the striatopallidal and hyperdirect loops are pushed toward higher frequencies by increasing G_STN–Proto_ (Figs. 6Bab and 7Bb). In this scenario, the oscillatory activity contains several modes (different frequencies expressed) and the pattern of phase-shift varies between modes. In the third scenario, the two oscillation-capable circuits interfere destructively, suppressing sustained oscillatory activity and stabilizing the network in the steady state. This is illustrated by the absence of sustained oscillations at striatopallidal synaptic weights sufficient to drive oscillations in isolation, but not when G_STN–Proto_ and G_GPi–Th_ are both non-zero (Figs. 6Bc and 7Bc). Across the full parameter space, certain combinations of generators gave rise to oscillations with a dominant frequency in the beta range in the Proto population. In the rat model, frequencies in the 15–20 Hz range emerged through the interaction of the striatopallidal loops with either the Proto or STN loop, although the dynamics were not straightforward (Fig 6A, top row). Specifically, low-beta oscillations generated by the striatopallidal loops in isolation, appearing around G_D2–Proto_ = 1.5, were suppressed by increasing either G_Proto–Proto_ or G_STN–Proto_, but could be restored by further increasing G_D2–Proto_, rescuing stable oscillations at slightly higher frequencies (Fig 6Ba). The relative phases between the populations in this region are close to the ones observed for oscillations driven by the striatopallidal loop only (Fig 4C). This rescue persisted until the Proto and STN loops became dominant, at which point they imposed high-frequency oscillations (> 50 Hz) that overwhelmed any beta-range activity. This dynamic was further disrupted by engagement of the hyperdirect loop through increases in G_GPi–Th_: the gradual frequency shift into the 15–20 Hz range was replaced by a sharp transition into a 25–30 Hz oscillatory regime (Fig 6Bb). These dynamics were also apparent in the STN power spectrum (Fig S2).

**Figure 6.**
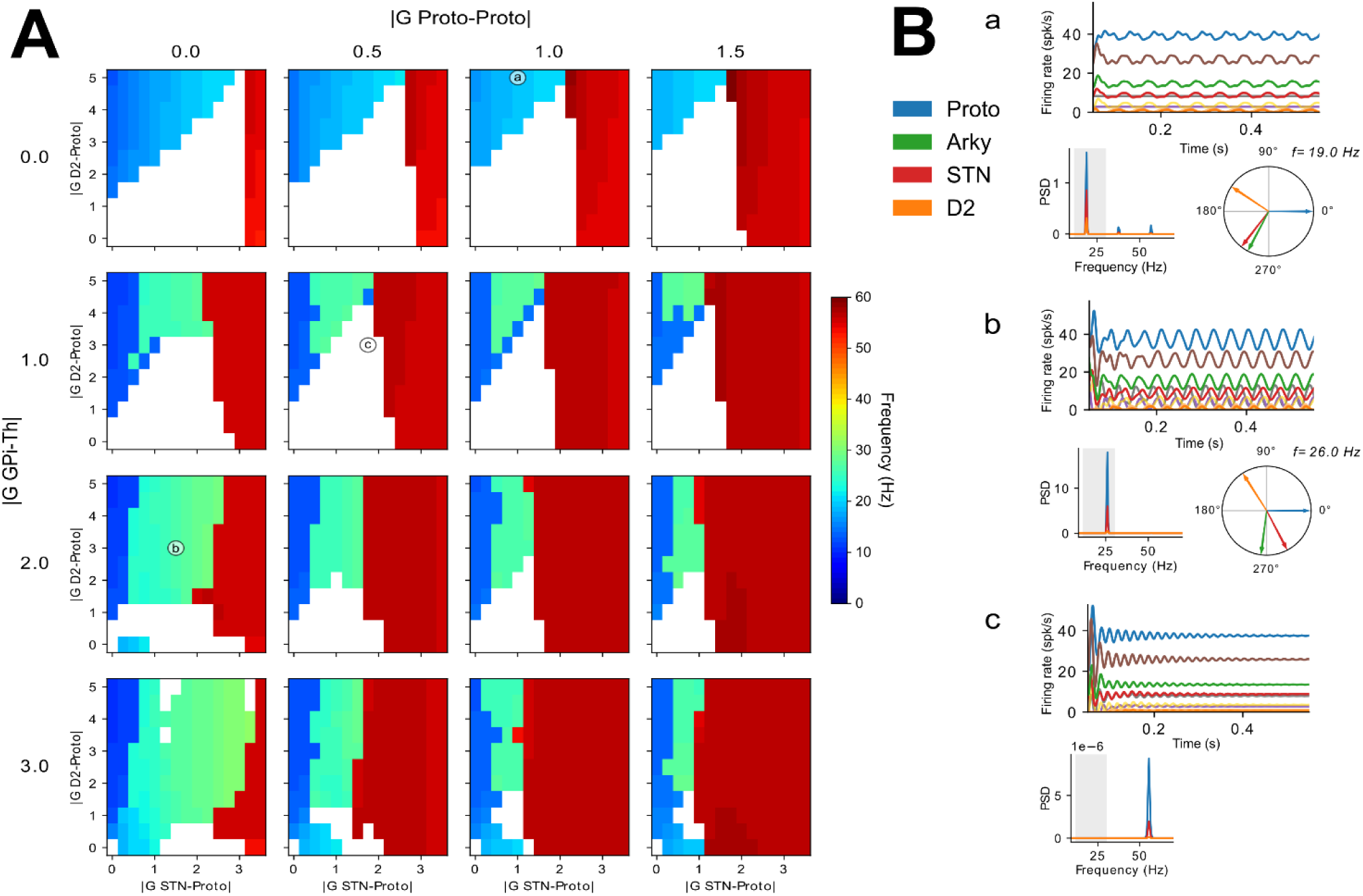
Oscillatory population dynamics under varying relative strengths of oscillation generators in the rate model of the rat BGTC network. A: Frequencies obtained in the Proto population by varying four synaptic weights involved in different generators: G_Proto–Proto_ is increased along the outer x axis, G_STN–Proto_ along the inner x axes, G_D2–Proto_ along the inner y axes, and G_GPi–Th_ along the outer y axis. White cells represent regions where oscillations are not stable (as in the example shown in C); colored cells represent the dominant frequency of stable oscillations in the Proto population, as determined by the peak in power spectral density of the mean activity of Proto neurons. B: Population dynamics generated at each of the three highlighted points a, b, and c in (A). Reported for each point are the mean population activities (top); the power spectral density of Proto, STN and D2 populations, with the light grey band marking the beta range (bottom left); and if oscillations are stable, the peak phase shift of Proto, Arky, STN and D2 populations relative to Proto at the dominant frequency (bottom right).

**Figure 7.**
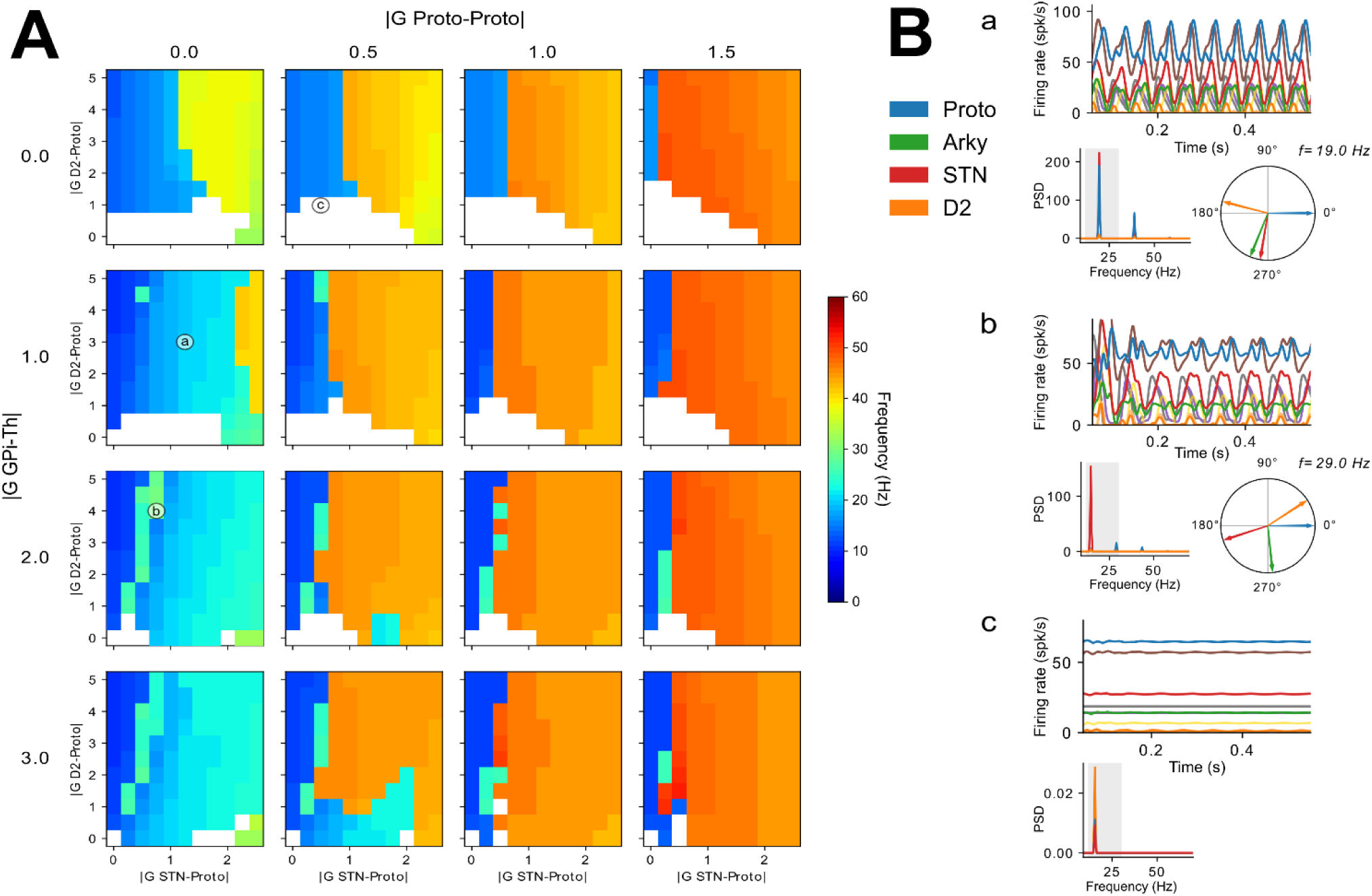
Oscillatory population dynamics under varying relative strengths of oscillation generators in the rate model of the monkey BGTC network. A: Frequencies obtained in the Proto population by varying four synaptic weights involved in different generators: G_Proto–Proto_ is increased along the outer x axis, G_STN–Proto_ along the inner x axes, G_D2–Proto_ along the inner y axes, and G_GPi–Th_ along the outer y axis. White cells represent unstable oscillations; colored cells represent the dominant frequency of stable oscillations in the Proto population. B: Population dynamics generated at each of the three highlighted points a, b, and c in (A). Reported for each point are the mean population activities (top); the power spectral density of Proto, STN and D2 populations, with the light grey band marking the beta range (bottom left); and if oscillations are stable, the peak phase shift of Proto, Arky, STN and D2 populations relative to Proto at the dominant frequency (bottom right).

The results from the monkey model (Fig. 7) were broadly consistent with those of the rat model, though the beta-frequency region was restricted due to the lower *Glim* thresholds of the Proto and STN loops. As a result, a larger proportion of simulations exhibited dominant frequencies above 40 Hz. A key difference appeared in the interaction between the striatopallidal and high-frequency loops (Fig. 7A, top row): while destructive interference in the rat model led to a steady asynchronous regime, the monkey model sustained stable oscillations at ∼15 Hz. The emergence of 25–30 Hz dominant frequencies also differed: rather than a sharp transition, it arose as a gradual shift from the low-beta range, jointly driven by the hyperdirect and STN loops (Fig. 7A, left column). Additionally, a narrow set of parameter combinations produced ∼28 Hz dominant frequencies outside this continuous shift (e.g., point b in Fig. 7A). However, when examining the STN power spectrum, these peak frequencies fell within the regular beta range (Fig. S3). The lowest observed frequencies (10–13 Hz) were generated by increased cortical feedback (non-zero G_GPi–Th_) and minimal STN loop involvement (low G_STN–Proto_). While the hyperdirect loop alone could drive these low-frequency oscillations, the striatopallidal loops could also contribute—when the D2-Proto synaptic gain was increased (Fig. 7A). To confront these findings with experimental evidence in NHPs (Tachibana et al., 2011) we tested the effect of pharmacological GPe inhibition on network activity when oscillations were driven solely by the hyperdirect loop. Despite the GPe not being part of the hyperdirect loop itself, partial inhibition of the GPe (70% blockade of Proto and Arkypallidal neurons) completely suppressed the ongoing 13 Hz oscillations, shifting the network to a steady non-oscillatory state (Fig. S4). This suppression occurred because reduced inhibitory input from Proto neurons to the STN led to disinhibition of STN neurons, which in turn increased GPi firing. The resulting inhibition of thalamic and cortical activity disrupted the hyperdirect feedback loop, precluding any oscillatory activity. Critically, this demonstrates that a nucleus can be necessary for sustaining oscillations without being part of the loop that generates them. Thus, the set of nuclei required for oscillations extends beyond the generator loop itself.

## Discussion

We investigated the spectral features and coherence patterns of synchronized beta-range oscillatory activity in models of the rodent and primate BGTC network strongly constrained by available anatomical and physiological data. Our approach combined analytical derivations of isolated sub-circuits with simulations of full dynamics in rate and spiking networks. Specific BGTC sub-circuits spontaneously generate synchronous low-beta oscillations in both species, such as striatopallidal loops (with Proto, D2, and FSI or Arky) and hyperdirect loop (cortex-STN-GPi-thalamus). While these sub-circuits oscillate at similar frequencies, phase angle distribution analyses across nuclei revealed unique patterns tied to each generator. This highlights the potential of multi-region phase relationships as an experimentally-accessible proxy for oscillatory mechanisms. Simulations also showed that interactions between generators can suppress oscillations via destructive interference or produce frequency shifts, including continuous transitions between low- and high-beta ranges or abrupt jumps (e.g., frequency doubling from loop interference). These interactions expand the achievable frequency range beyond individual generators.

Many computational studies have long explored beta oscillations in BG models (Pavlides et al., 2015; Rubin, 2017). Early research identified the STN–GPe recurrent loop as a potential generator of oscillations within the BG (Plenz and Kital, 1999; Terman et al., 2002). Given the prominence of beta-band activity in both the STN and GPe and their relevance to DBS (Krack et al., 2003; Vitek et al., 2012), this loop was proposed to drive pathological oscillations. This hypothesis is supported by multiple modeling studies demonstrating that the subthalamopallidal loop oscillates within the beta range (Nevado-Holgado et al., 2010; Kumar et al., 2011; Park et al., 2011; Merrison-Hort et al., 2013; Pasillas-Lépine, 2013; Pavlides et al., 2015; Shouno et al., 2017; Koelman and Lowery, 2019; Chen et al., 2020). However, these studies rely on slow neuronal and/or synaptic time constant in the STN–GPe interactions to achieve beta-range frequencies, while both *in vitro* and *in vivo* experimental evidence reveals fast transmission between the two nuclei (Kita et al., 1983; Kita and Kitai, 1991; Nambu et al., 2000; Baufreton et al., 2005, 2009; Chu et al., 2015). Using experimentally derived time constants, we showed that STN–GPe oscillations fall above the beta range in both rat and monkey models, making it unlikely to be the sole or main driver of pathological beta activity. Consistent with our findings, other studies have proposed that the hyperdirect loop drives abnormal low-beta oscillations upon dopamine depletion (Leblois et al., 2006; Pavlides et al., 2015) and that pallidostriatal feedback promotes beta synchrony (Corbit et al., 2016; Ortone et al., 2023; Azizpour Lindi et al., 2024; Zang et al., 2024). Generator interactions could also suppress oscillations through destructive interference. In noisy systems, this may manifest as transitions between transient and sustained oscillations, paralleling the shift from physiological to pathological beta (Feingold et al., 2015; Deffains et al., 2018; Duchet et al., 2021). Dopamine loss may push transient bursts toward sustained oscillations by altering synaptic gains in critical BGTC pathways.

Data from 6-OHDA rats suggest the hyperdirect loop is not necessary for abnormal beta oscillations, as neither cortex nor STN is required for their expression (De la Crompe et al., 2020). Instead, evidence supports a central role for the striatopallidal circuit (Sharott et al., 2017; De la Crompe et al., 2020, 2025). Oscillations in rats typically occur in the high-beta range (15–30 Hz) (Mallet et al., 2008b; Avila et al., 2010), with frequencies shifting based on behavioral state: ∼20 Hz under anesthesia (Mallet et al., 2008a; West et al., 2018), accelerating to 25–30 Hz at rest and >30 Hz during locomotion in the cortex (Degos et al., 2009), STN (Lehmkuhle et al., 2009; Jávor-Duray et al., 2015) and SNr (Avila et al., 2010; Brazhnik et al., 2014; Jiang et al., 2019). This shift in frequency resemble our model’s frequency changes when varying generator strengths, suggesting pathological beta in rats emerges from multiple generators whose relative strengths depend on cortical state. In our simulations, 15–20 Hz frequencies arose from striatopallidal loops entrained by the subthalamopallidal loop or by Proto lateral inhibition, the latter high-frequency generators shifting frequencies upward. For 20–30 Hz, interactions involved subthalamopallidal, striatopallidal, and hyperdirect loops. The Proto loop mechanism aligns with De la Crompe et al. (2020), where STN lesions did not suppress beta oscillations, suggesting that the subthalamopallidal loop is not a necessary high-frequency generator in this context. Phase shifts between recorded populations further supported this scenario. Whether higher frequencies in awake rats depend on STN integrity and the relative phases of populations remains untested.

MPTP-induced monkeys exhibit beta oscillations in the low-beta/alpha ranges (8–15 Hz) (Raz et al., 2001; Wichmann and DeLong, 2003; Leblois et al., 2007; Deffains and Bergman, 2019). In our monkey model, the striatopallidal and hyperdirect loops spontaneously oscillated in the low-beta range, while the STN–GPe loop oscillated above the beta band. In simulations involving multiple generators, low-beta oscillations persisted only when cortical feedback was present, implicating the hyperdirect loop as the key driver. This aligns with findings by Tachibana et al. (2011), who demonstrated that STN inputs are critical for sustaining oscillations. Low-beta oscillations were most robust when the STN–GPe loop had minimal involvement, effectively ruling it out as a meaningful contributor to their generation. Nevertheless, the GPe, through its inhibitory projections to the STN and GPi, is necessary to sustain the oscillations driven by the hyperdirect loop, as the firing rate changes induced by GPe silencing preclude oscillatory activity. This is consistent with Tachibana et al. (2011), who also identified the GPe as essential for oscillations.

Phase relationships can distinguish between competing beta-generation hypotheses and could serve as a biomarker for the circuit underlying pathological oscillations in PD. These relationships are conserved for multiple interacting generators as long as one oscillator dominates, but become multimodal phase distributions when several frequencies peaks are observed. In 6-OHDA rats, the reported antiphase relationship between Proto/STN and Proto/Arky neurons, and in-phase between STN/Arky (Mallet et al., 2008a, 2012; De la Crompe et al., 2020) aligns most closely with the striatopallidal generator (Mallet et al., 2008a; West et al., 2018; De la Crompe et al., 2020; Azizpour Lindi et al., 2024). In NHPs, phase analyses are limited due to a lack of simultaneous multi-population recordings. In PD patients, motor cortex–STN/GPi coherence shows cortical activity leading STN/GPi by ∼20 ms for <30 Hz oscillations (Marsden et al., 2001; Williams et al., 2002), consistent with hyperdirect-loop generation. It should be noted that neuronal model choice (e.g., LIF vs. quadratic integrate-and-fire) can alter phase relationships in the STN–Proto network (Tse et al., 2026), suggesting model-dependent predictions. Further work is needed to clarify phase–spike relationships in realistic networks and characterize spike generation in BG neurons across species.

A key limitation is the scarcity of experimental synaptic and cellular parameters for the monkey model, given the scarcity of in vitro NHP studies, restricting us to rate models. Spiking networks capture short-timescale dynamics but lack reliable parameters estimation from NHP studies. Our rat LIF spiking model matched the rate model’s frequency ranges and mean phases, with broader phase distributions consistent with experiments. Rate models suit beta’s slow timescales but may underestimate *in vivo* variability (e.g., phase distributions). The absence of spike-frequency adaptation in rate models may also explain discrepancies in fast dynamics (e.g., STN inhibition latency). Incorporating cellular-level detail, as in spiking models, could refine predictions, but this remains constrained by data availability in NHPs.

The BGTC model included most major synaptic projections, although some known connections were omitted for the sake of simplification. We minimized Arky population projections due to shared architecture between rat and monkey models. The presence of arkypallidal neurons in monkeys GPe is debated: while emerging evidence points to two distinct GPe subpopulations that may be homologous to the Proto and Arky classes in rodents (Katabi et al., 2023), this remains uncertain. We retained only Proto-to-Arky input, omitting weaker D2/STN inputs to Arky (Aristieta et al., 2021). Striatal MSN collaterals (within/between D1/D2) were excluded, as prior work showed their removal has little effect on striatal firing balance (Damodaran et al., 2014), unlike FSI–MSN projections. Some approaches even omit D1 neurons entirely due to their low firing rates under PD (Zang et al., 2024). Finally, synaptic gains are a central free parameter in rate models and lack direct experimental measurement. Combining optogenetics, electrophysiology, and computational inference could estimate these gains experimentally *in vivo*, improving predictions of generator dominance and interactions. Such experiments could also reveal how dopamine depletion and progressive PD pathology alter the various synaptic gains in the network, particularly the balance between the oscillation-generating circuits presented here.

## Supplemental Material

**Table S1.** Population sizes. Values were estimated from the experimental literature and used to compute the K_sim_ connectivity values for the models (see Eq. 1). NA: not available.

| Population | Values in the rat brain | Values in the monkey brain |
| --- | --- | --- |
| Striatum | 2,790,000 (Oorschot, 1996) | 30,400,000 (only MSNs) (Yelnik et al., 1991) |
| GPe | 46,000 (Oorschot, 1996) | 502,000 (Hardman et al., 2002) |
| Arky | 11,500 (25% of GPe) (Mallet et al., 2012) | 65,260 (13% of GPe) (Katabi et al., 2023) |
| Cortex | 17,000,000 (Zheng and Wilson, 2002) | 100,000,000 (Bar-Gad et al., 2003) |
| D1 | 1,325,250 (47.5% of striatum) (Oorschot, 1996) | 15,200,000 (50% of striatum) (Yelnik et al., 1991) |
| D2 | 1,325,250 (47.5% of striatum) (Oorschot, 1996) | 15,200,000 (50% of striatum) (Yelnik et al., 1991) |
| FSI | 111,600 (4% of striatum) (Tepper et al., 2010) | 1,520,000 (5% of striatum) (Yelnik et al., 1991) |
| GPi | 3,200 (Oorschot, 1996) | 156,000 (Hardman et al., 2002) |
| Proto | 32,200 (70% of GPe) (Mallet et al., 2012) | 436,740 (87% of GPe) (Katabi et al., 2023) |
| STN | 13,560 (Oorschot, 1996) | 94000 (Hardman et al., 2002) |
| Thalamus | NA | NA |

**Table S2.**
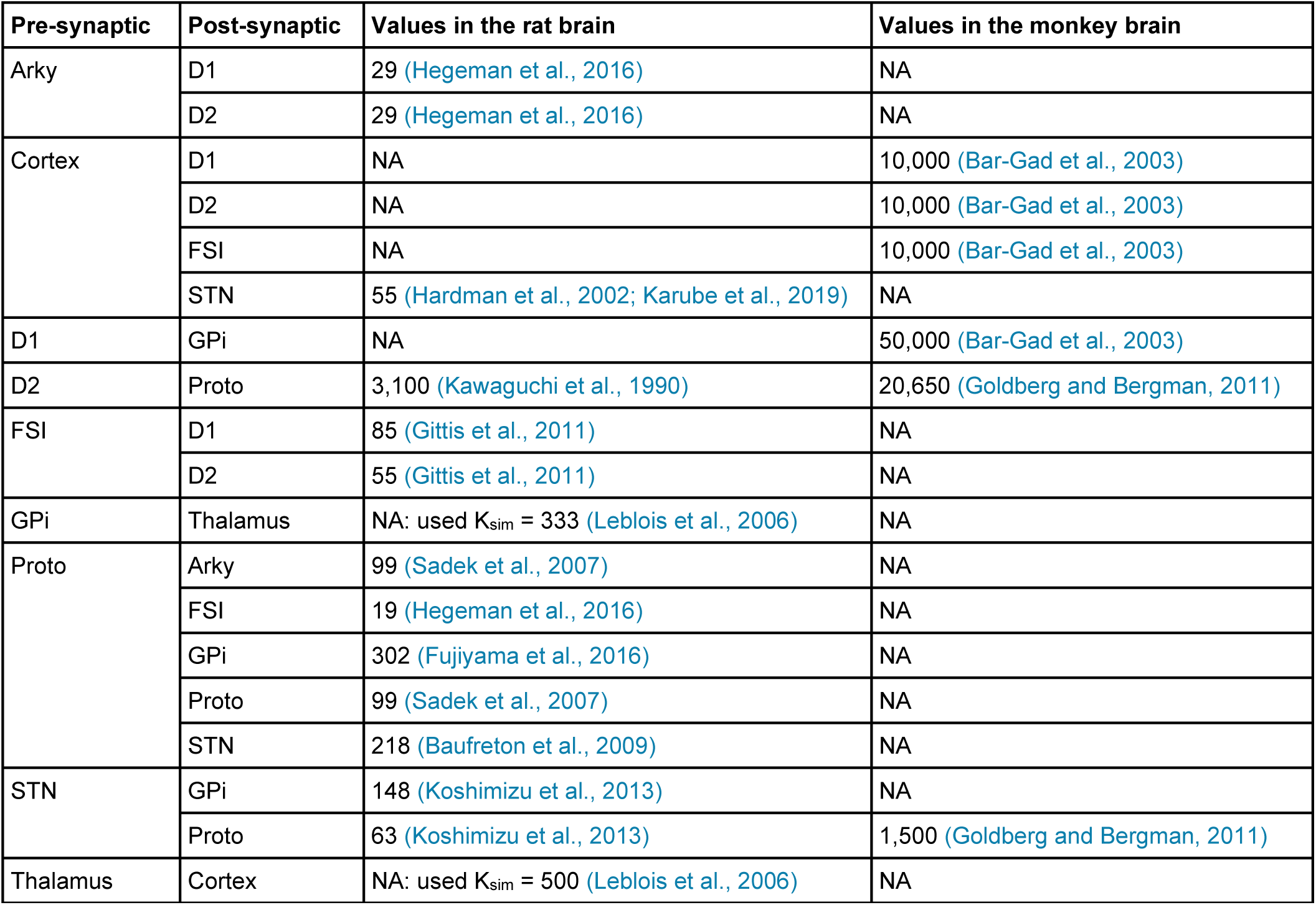
Number of presynaptic neurons converging onto a postsynaptic neuron. Values were estimated from the experimental literature and used to compute the K_sim_ connectivity values for the models (see Eq. 1). When values were not available (NA), the final K_sim_ value used in the model was generalized from the other species model. When values were not available in either rodents nor monkeys, we used K_sim_ values from the computational literature directly.

| Pre-synaptic | Post-synaptic | Values in the rat brain | Values in the monkey brain |
| --- | --- | --- | --- |
| Arky | D1 | 29 (Hegeman et al., 2016) | NA |
|  | D2 | 29 (Hegeman et al., 2016) | NA |
| Cortex | D1 | NA | 10,000 (Bar-Gad et al., 2003) |
|  | D2 | NA | 10,000 (Bar-Gad et al., 2003) |
|  | FSI | NA | 10,000 (Bar-Gad et al., 2003) |
|  | STN | 55 (Hardman et al., 2002; Karube et al., 2019) | NA |
| D1 | GPi | NA | 50,000 (Bar-Gad et al., 2003) |
| D2 | Proto | 3,100 (Kawaguchi et al., 1990) | 20,650 (Goldberg and Bergman, 2011) |
| FSI | D1 | 85 (Gittis et al., 2011) | NA |
|  | D2 | 55 (Gittis et al., 2011) | NA |
| GPi | Thalamus | NA: used $K_{sim} = 333$ (Leblois et al., 2006) | NA |
| Proto | Arky | 99 (Sadek et al., 2007) | NA |
|  | FSI | 19 (Hegeman et al., 2016) | NA |
|  | GPi | 302 (Fujiyama et al., 2016) | NA |
|  | Proto | 99 (Sadek et al., 2007) | NA |
|  | STN | 218 (Baufreton et al., 2009) | NA |
| STN | GPi | 148 (Koshimizu et al., 2013) | NA |
|  | Proto | 63 (Koshimizu et al., 2013) | 1,500 (Goldberg and Bergman, 2011) |
| Thalamus | Cortex | NA: used $K_{sim} = 500$ (Leblois et al., 2006) | NA |

**Table S3.**
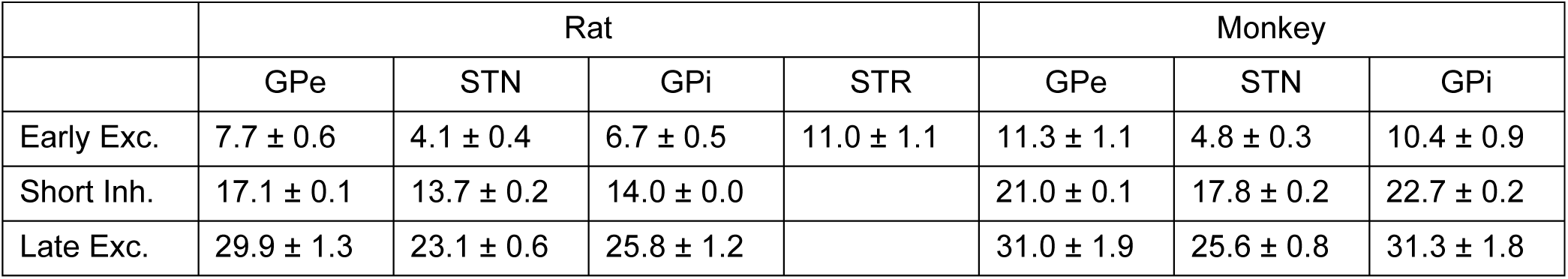
Response latencies to cortical stimulation in the two rate models.

**Figure S1.**
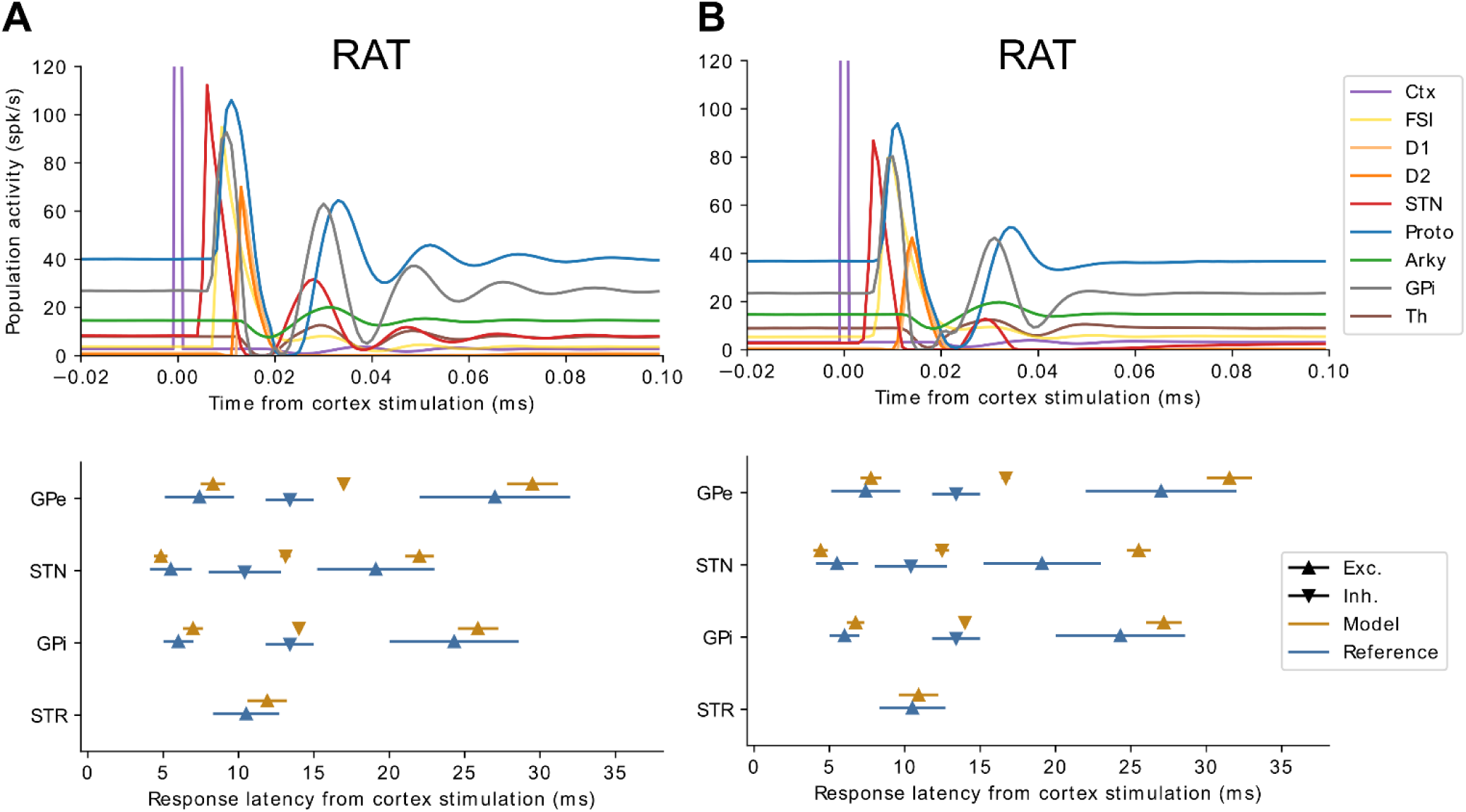
Effect of cortical stimulation with alternative parameters for the rat model. Average population activities (top) and mean and s.d. of response latencies (bottom). **A.** Results of a simulation with Δ_Proto–STN_ = 0.5 ms. **B.** Results of a simulation with STN spike frequency adaptation implemented as an additional adaptation current *A_adapt_* such that 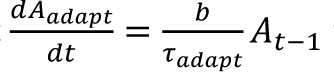 with *b* = 60, *τ_adapt_* = 50 ms, and *A* the instantaneous activity.

**Figure S2.**
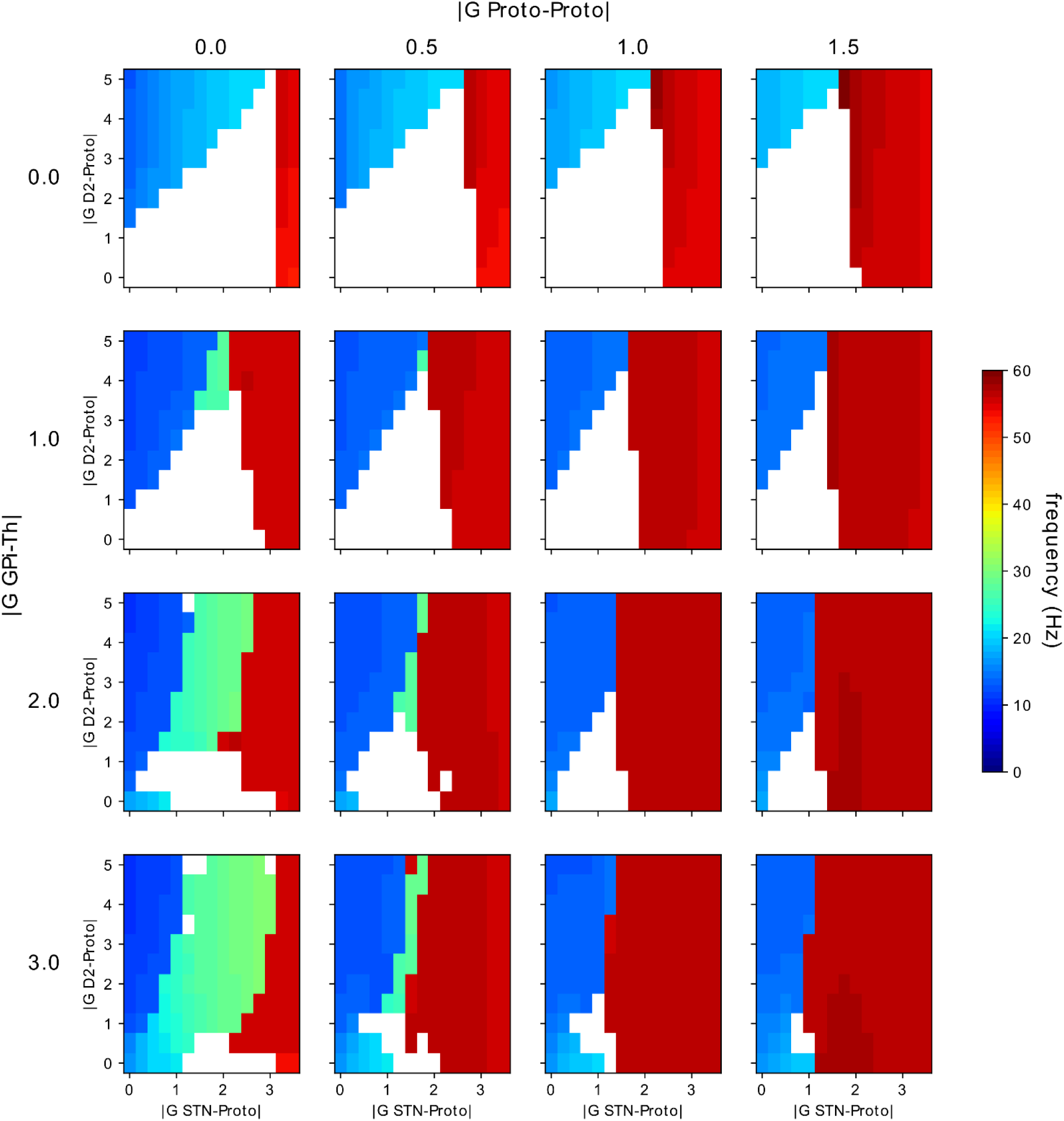
Effect of varying synaptic gains of the rat model on the peak frequency of the STN population. White cells represent unstable oscillations, colored cells represent the dominant frequency of stable oscillations in the STN population.

**Figure S3.**
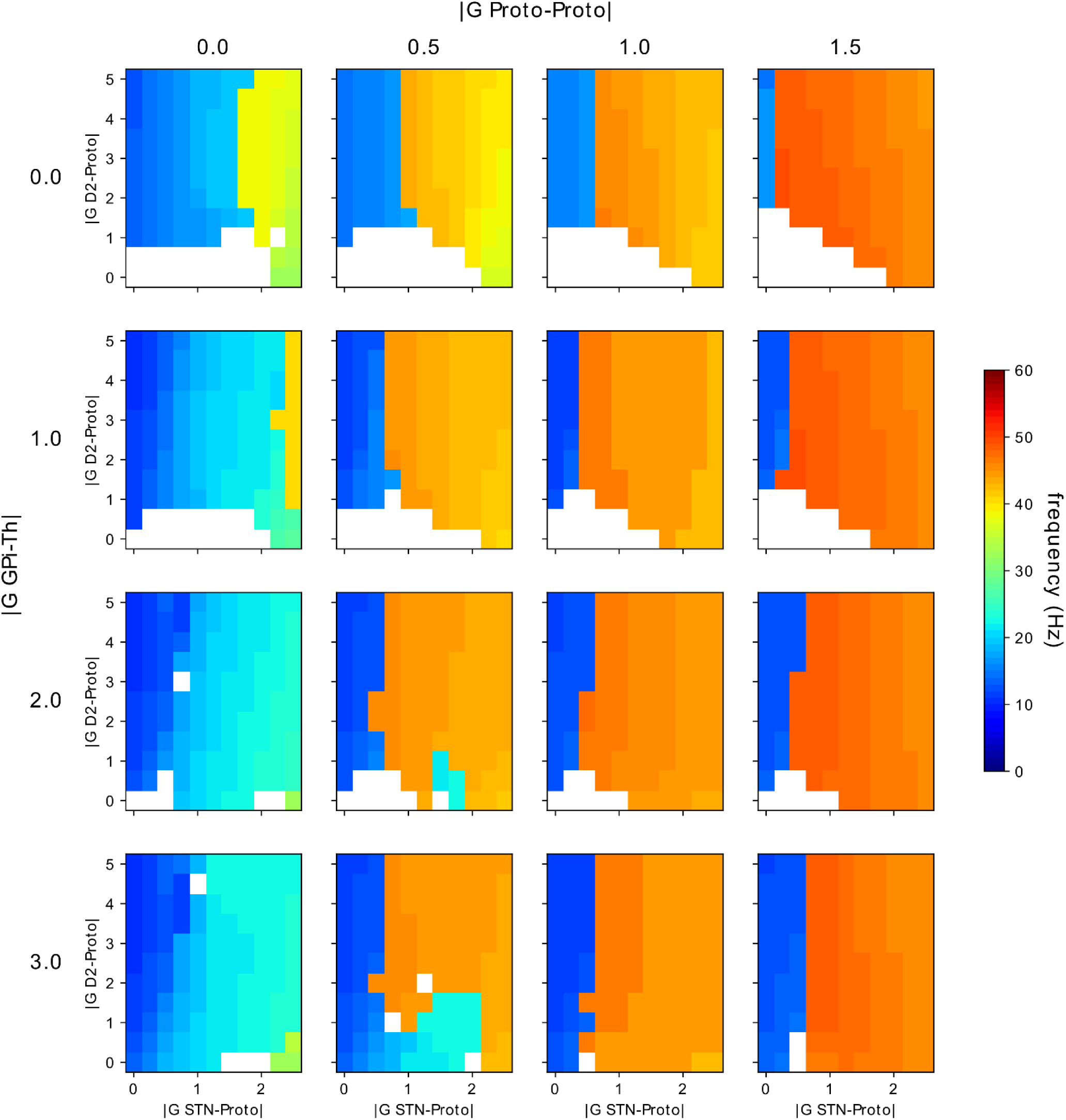
Effect of varying synaptic gains of the monkey model on the peak frequency of the STN population. White cells represent unstable oscillations, colored cells represent the dominant frequency of stable oscillations in the STN population.

**Figure S4.**
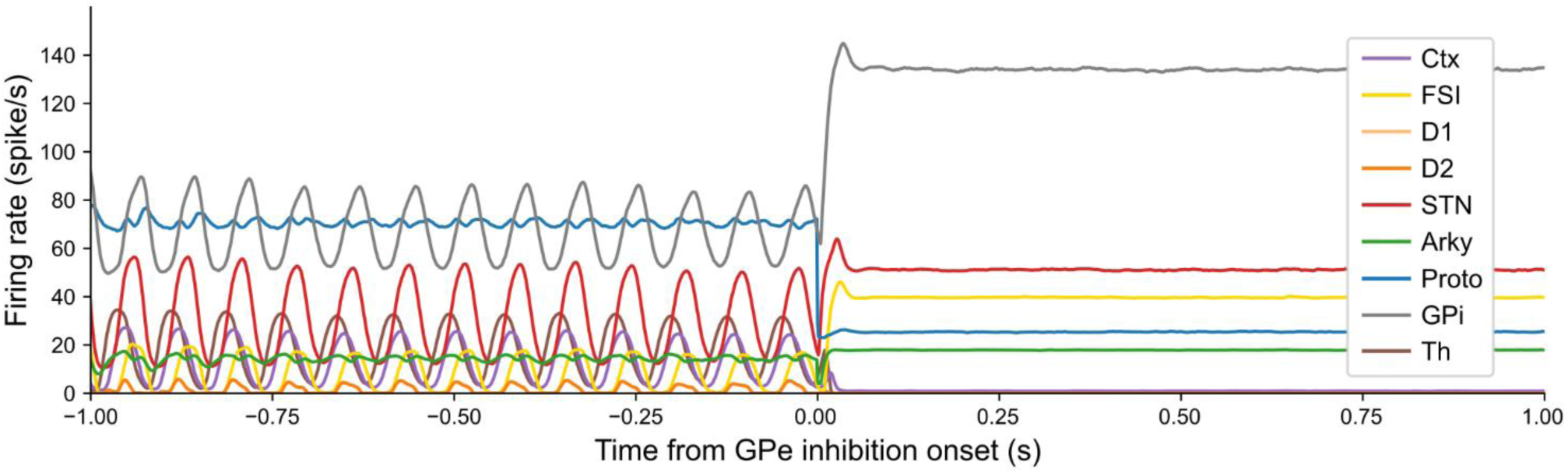
Oscillations driven by the hyperdirect loop in the monkey rate model are interrupted by GPe silencing. The network is set with G_Proto–Proto_ = 0, G_STN–Proto_ = 0.25, G_D2–Proto_ = –2, G_Ctx–STN_ = 2, and all other weights set to 1. Introducing inhibition of the GPe populations disinhibits STN and increases GPi firing rate, silencing Th and Ctx and abolishing stable oscillations.

